# Assembly and substrate engagement mechanism of the bacterial proteasome activator Bpa

**DOI:** 10.1101/2025.10.01.679806

**Authors:** Bradley T. V. Davis, Enrico Rennella, Anisha Haris, Jakub Ujma, David Bruton, Keith Richardson, Kevin Giles, Lewis E. Kay, Siavash Vahidi

**Author notes:** These authors contributed equally (Bradley T. V. Davis, Enrico Rennella). Correspondence to: Siavash Vahidi; or Lewis E. Kay.

## Abstract

The <u>b</u>acterial <u>p</u>roteasomal <u>a</u>ctivator Bpa (Rv3780) is an ATP-independent regulatory particle of the *Mycobacterium tuberculosis* proteasome system. Bpa recruits substrates as a dodecamer and triggers the gate opening of the proteasome 20S core particle; however, the structural basis for its oligomerization and substrate recognition remains unclear. Here, we define the temperature-sensitive oligomerization mechanism of Bpa and elucidate its interaction with a non-native substrate. Using size-exclusion chromatography, charge detection mass spectrometry, and pulsed hydrogen/deuterium exchange mass spectrometry (HDX-MS), we show that Bpa reversibly assembles into a dodecameric ring from dimeric and tetrameric species in a temperature-dependent manner. We used HDX-MS to map the oligomerization interfaces during Bpa assembly. Methyl transverse relaxation optimized spectroscopy (TROSY)-based NMR experiments and site-specific truncations further validate the existence of discrete tetrameric and dodecameric states. To overcome the limitations posed by the poor solubility of the native substrates of Bpa, we establish the DNA-binding domain of hTRF1 as a surrogate substrate. Bpa binds hTRF1 and mediates its degradation in a 20S CP-dependent manner. We quantify the affinity and stoichiometry of the Bpa-hTRF1 interaction using methyl-TROSY NMR, identifying a 12 Bpa subunit : 3 hTRF1 binding ratio with micromolar affinity that is modulated by salt concentration. Our NMR-based mapping experiments pinpoint the interaction surfaces on both Bpa and hTRF1, revealing key hydrophobic residues that mediate substrate engagement. This work uncovers a thermosensitive switch regulating Bpa oligomerization and activity and introduces a tractable substrate for dissecting proteasomal recognition in *M. tuberculosis*.

## Introduction

*Mycobacterium tuberculosis* (*Mtb*) uses a rare prokaryotic proteasome system to resist nitrosative and oxidative toxicity in host macrophages^1,2^. Thus, the *Mtb* proteasome and its interacting partners are novel therapeutic targets for developing antibacterials against multidrug-resistant *Mtb*^3–5^. The *Mtb* 20S core particle (CP) consists of an assembly of homoheptameric rings, arranged in a α7-β7-β7-α7 architecture, that sequesters fourteen catalytic sites, one within each β-subunit (Fig. 1a). Proteasomal activity is tightly controlled by a gating mechanism within the α7-rings^6^. As a result, the 20S CP alone can only degrade small, unstructured substrates that are able to passively diffuse through the closed gate. Activation of the 20S CP requires the binding of unstructured C-terminal GQYL motifs of regulatory particles (RPs) to an α7-ring, triggering gate opening and enabling substrate entry^6–9^ (Fig. 1b). RPs also engage, unfold, and translocate substrates into the 20S CP for degradation. *Mtb* encodes two known RPs that interact with the 20S CP: Mpa, and Bpa^7,10–14^. The hexameric, ATP-dependent RP Mpa (<u>m</u>ycobacterial <u>p</u>roteasome <u>A</u>TPase; Rv2115c), unfolds and translocates pupylated substrates into the 20S CP for degradation. In contrast, the Bpa (<u>b</u>acterial <u>p</u>roteasome <u>a</u>ctivator; Rv3780) RP functions independently of the pupylation pathway and mediates the degradation of a distinct set of substrates in an ATP-independent fashion^7,12^. Bpa is essential for *Mtb* persistence under heat shock and oxidative stress by degrading, among other substrates^7,11^, HspR, a transcriptional repressor that regulates the Hsp70/40 chaperone system^2,12,15,16^. While Mpa’s substrate recruitment is well characterized^17–19^, the substrate recognition mechanism of Bpa remains poorly understood, with HspR the only well-established native substrate characterized to date^2,11,15^.

**Figure 1.**
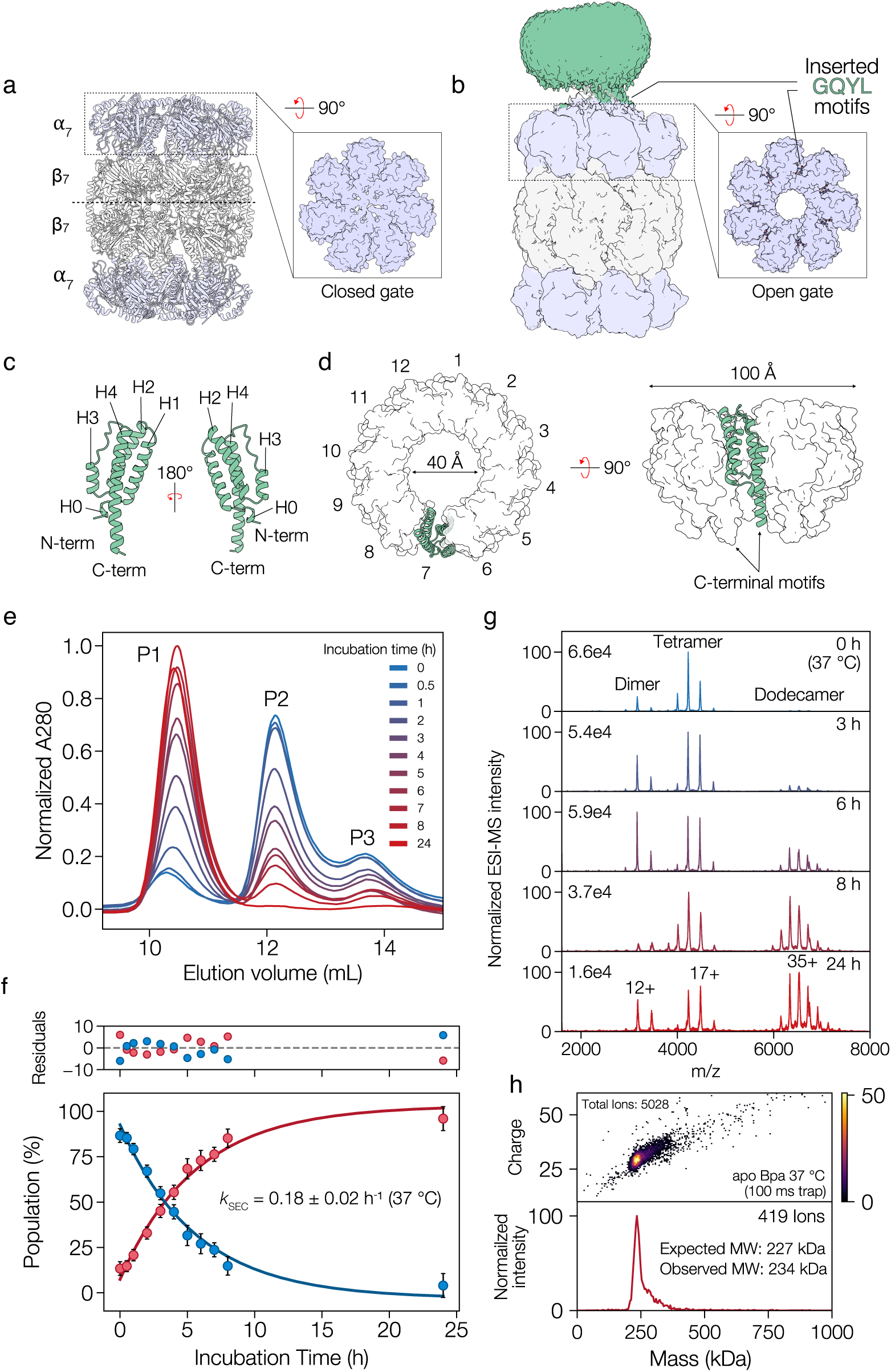
Bpa undergoes a temperature-dependent oligomerization. **a)** Cartoon representation of an unbound 20S CP (PDB: 9CE5)^48^. The expanded box depicts the closed gate conformation of the gating residues (PDB: 6BGO)^7^; (**b**) Density map of Bpa bound to the 20S CP (EMDB: 19150)^20^. The expanded box highlights the open-gate conformation induced by insertion of the Bpa C-terminal GQYL motif into a pocket formed at the interface between adjacent subunits of the α7-ring (PDB: 6BGO)^7^; (**c**) Cartoon representation of an individual Bpa subunit, highlighting the four-helix bundle fold and the flexible C-terminal region containing the ^171^GQYL^174^ motif. Residues 145-174 are not resolved in the crystal structure (PDB ID: 5LFJ)^13^; (**d**) Top and side view of an individual Bpa subunit (cartoon) within the 227.3 kDa homododecameric Bpa ring (surface); (**e**) SEC chromatograms indicating a shift in the Bpa oligomeric equilibrium as a function of incubation time (37 °C), after equilibration of Bpa at 4 °C. The large molecular weight (MW) species is denoted as P1 (peak 1), while the two lower MW species are denoted as P2 and P3 (peaks 2 and 3); (**f**) Changes in the fractional populations of the P1 (red) and P2+P3 (blue) species from analysis using an exponential decay model to extract an association rate (*k*). Residuals of the decay model fitting are displayed in the plot above; (**g**) Nano-ESI native MS displays changes in the oligomeric equilibrium of Bpa upon incubation at 37 °C, starting from a distribution that was generated at 4 °C. The species in solution were identified as dodecameric (P1), tetrameric (P2), and dimeric (P3) Bpa oligomers. All spectra were normalized to the highest intensity peak in each spectrum, with these values given in the top left corner. The predominant charge state for each species is denoted on the 24 h spectrum; and (**h**) CD-MS analysis of Bpa incubated overnight at 37 °C shows that WT Bpa predominantly adopts a dodecameric state.

Elegant structural studies have elucidated the architecture of substrate-free (apo) Bpa^12^ and its binding mechanism to the 20S CP^13,20^. Each Bpa subunit is comprised of a four-helix bundle that oligomerizes into a 228-kDa homododecameric ring^11,12^ (Fig. 1c, d). Two fundamental aspects of Bpa function remain poorly understood. First, the mechanism by which Bpa assembles into its dodecameric form is unknown, and it is unclear whether this assembly is constitutive or regulated by environmental cues. Second, the molecular basis by which Bpa recognizes and binds its substrates remains incompletely understood. von Rosen et al.^2^ used mutagenesis and biochemical assays in studies involving HspR, to show that Bpa recognizes and binds HspR through a bipartite mechanism, engaging a folded domain for stable association, while relying on a disordered, sequence-specific C-terminal tail to initiate translocation and degradation by the proteasome. This indicates, that at least for this substrates, intrinsic disorder alone is insufficient for Bpa recruitment, and that both structural and sequence features are critical for substrate selection. In subsequent work^20^, the same group used cross-linking mass spectrometry and electron cryomicroscopy (cryo-EM) of chemically-crosslinked Bpa-HspR complexes to implicate Bpa residues H131, F138, and R145 in substrate binding. Substantial heterogeneity in Bpa-HspR complexes precluded high-resolution structure determination, leaving the precise binding interface between Bpa and its substrates unresolved.

Here, using a combination of mass spectrometry (MS)- and nuclear magnetic resonance (NMR) spectroscopy-based tools, we map the dynamic interfaces that mediate Bpa oligomerization and substrate recognition. By leveraging a non-native model substrate, we show that Bpa recognizes short hydrophobic motifs embedded within disordered regions of substrates. This interaction is mediated through solvent-accessible hydrophobic bands on the inner surface of the Bpa dodecameric ring, is modulated by ionic strength, and exhibits defined stoichiometry. Our work supports a model in which thermally regulated Bpa assembly enables conditional proteasomal activation in response to proteotoxic stress and offers broader implications for understanding regulated proteolysis across mycobacteria.

## Results

### Bpa undergoes temperature-dependent oligomerization

We heterologously expressed and purified wild-type (WT) *Mtb* Bpa. In the process of purification optimization, we observed a striking temperature-dependent shift in elution profiles during the final size exclusion chromatography (SEC) step. Samples incubated at 4 °C displayed a different chromatographic peak distribution compared to those incubated at 37 °C, consistent with changes in particle size. To quantify this process, we incubated Bpa at 37 °C following extended storage at 4 °C and analyzed Bpa particle size at defined time intervals by SEC. Chromatograms revealed three distinct peaks corresponding to a high-molecular weight assembly (P1) and two lower-molecular weight species (P2 and P3), whose relative abundances changed with incubation time (Fig. 1e). At the onset of incubation at 37 °C, the chromatograms were dominated by the smaller P2 and P3 species. These species progressively converted to the larger P1 species over time. We quantified this transition by integrating each peak and fitting the increase in P1 abundance using a first-order exponential model, yielding an apparent rate constant *k*SEC of 0.18 ± 0.02 h^-1^ (Fig. 1f). Importantly, Bpa oligomerization is fully reversible with respect to temperature changes, consistent with a dynamic assembly equilibrium.

To determine the absolute molecular weights of the Bpa oligomeric states, we used size exclusion chromatography coupled to multi-angle light scattering (SEC-MALS). The high-molecular weight species (P1) corresponded to the expected 228 kDa-dodecameric Bpa complex (Fig. S1). The lower-molecular weight species eluting at ∼18 mL exhibited a broad asymmetric peak indicative of oligomeric heterogeneity, precluding accurate molecular weight determination by MALS (Fig. S1).

We used native electrospray ionization mass spectrometry (native MS) to monitor the Bpa assembly process. We first used a quadrupole time-of-flight (Q-TOF) mass spectrometer equipped with a nano-electrospray source to determine the exact molecular weights of all species. Bpa samples were incubated at 4 °C for 24 hours, then transferred to 37 °C to trigger assembly. Aliquots were subsequently taken at defined time points for native MS measurement. The native mass spectra display three sets of charge state distributions corresponding to dimeric, tetrameric, and dodecameric Bpa (Fig. 1g). The relative intensity of the dodecameric species increased with incubation time at 37 °C, corroborating our SEC data (Fig. 1e). Notably, even after overnight incubation, low levels of dimeric and tetrameric species were still detected in our spectra, features absent from our SEC data (Fig. 1e). We attribute this apparent discrepancy to the enhanced transmission and detection efficiency of lower mass ions on our Q-TOF instrument^21,22^.

To mitigate mass-dependent detection bias and to accurately identify the final distribution of oligomeric states, we turned to charge detection mass spectrometry (CD-MS). We used a prototype CD-MS instrument equipped with an electrostatic linear ion trap (ELIT) developed by the Waters Corporation. CD-MS measures the charge (z) and mass-to-charge ratio (m/z) of individual ions as they oscillate within an ELIT device. By directly detecting the image current produced by each ion, CD-MS overcomes the limitations of conventional MS for analyzing highly heterogeneous or high-mass ions, where charge state resolution is often compromised. This single-ion approach enables accurate mass determination of large noncovalent complexes regardless of peak overlap or mass distribution complexity. We analyzed Bpa samples incubated at 37 °C with an ion trapping time of 100 ms. The resulting two-dimensional mass versus charge and one-dimensional mass histograms displayed a single dominant population corresponding to fully assembled dodecameric Bpa, with negligible signal from smaller oligomers (Fig. 1h). The absence of signal from dimers and tetramers, even under low ion trapping conditions that favour detection of small analytes, independently verifies our SEC measurements that Bpa fully assembles into dodecamers upon incubation at 37 °C.

### H/D exchange mass spectrometry probes the oligomerization interface of Bpa

We used pulsed hydrogen/deuterium exchange mass spectrometry (HDX-MS) to define the structural interfaces that mediate the oligomerization of Bpa. HDX-MS is a powerful technique for probing conformational dynamics and protein-protein interactions in solution^23,24^. HDX is governed by the rate at which backbone amide hydrogens exchange with deuterium in a D2O-based buffer, a process modulated by solvent accessibility and hydrogen bonding networks^24,25^. Regions buried within oligomerization interfaces or those stabilized by secondary structure exhibit slower deuterium uptake, whereas dynamic or solvent-exposed segments exchange more rapidly^26^. Importantly, conformational changes during assembly can either lead to an increase or a decrease in HDX rates, depending on the structural rearrangements involved^27,28^. Localization of these changes is achieved by proteolytic digestion under conditions that minimize back-exchange, followed by LC-MS analysis of deuterated peptides^29–31^.

We performed time-resolved pulsed HDX-MS to monitor changes in Bpa assembly in a spatially-resolved manner. Samples of unassembled Bpa (maintained at 4 °C) were incubated at 37 °C to initiate assembly. At defined time points, aliquots were removed and pulsed with D2O (pHcorr 7.0) for 10 seconds. These samples were next quenched to stop deuterium incorporation, and were flash-frozen and stored at -80 °C until LC-MS analysis. Our HDX-MS workflow yielded 87 peptides with quantifiable deuterium incorporation, covering 100% of the Bpa sequence with an average peptide redundancy level of 5.60 (Fig. S2, Table S2). The raw unprocessed spectra exhibited a mixture of unimodal and bimodal isotopic distributions, consistent with a heterogeneous conformational ensemble^32^.

As a starting point for interpreting our HDX-MS data, we computed differential deuterium uptake at each time point relative to the unassembled state and visualized these differences using a heat map (Fig. 2a). We also colour-coded the available structure of dodecameric Bpa using this heat map (Fig. 2b, c)^33^. This analysis revealed substantial shifts in deuterium uptake as a function of assembly time. Regions known to participate in the canonical dodecamer subunit interface, including helix H1, exhibited progressively reduced deuterium incorporation upon incubation at 37 °C, consistent with the burial of these surfaces during oligomerization. For example, peptides 40-53, 61-85, and 98-116, spanning major portions of H1, H2, and H4, respectively, showed marked protection from exchange over time (Fig. 2b, c), indicative of decreased solvent accessibility and reinforced hydrogen bonding due to subunit packing relative to the dimeric and tetrameric structures that predominate at *t* = 0 (Fig. 1e, g). In contrast, residues located in the flexible C-terminal H4 region displayed increased deuterium uptake upon dodecamer formation. We interpret this increase as evidence of disengagement from interactions that stabilize lower-order oligomers. This interpretation is consistent with expectations based on a previously-determined crystal structure (PDB: 5IEU) of a truncated Bpa variant (44-153) that revealed a tetramer stabilized by inter-subunit H4-H4 interactions^12^ that would protect backbone amides in these helical regions from exchange with solvent (Fig. S3). Notably, this interface is distinct from the subunit packing observed within the context of the dodecameric Bpa assembly. Our data, thus, indicate that as the temperature increases the oligomerization equilibrium shifts toward dodecamer formation, H4 disengages from tetramer-specific interfaces, so that this region becomes increasingly solvent-exposed and exchange-competent (Fig. 2b, c).

**Figure 2.**
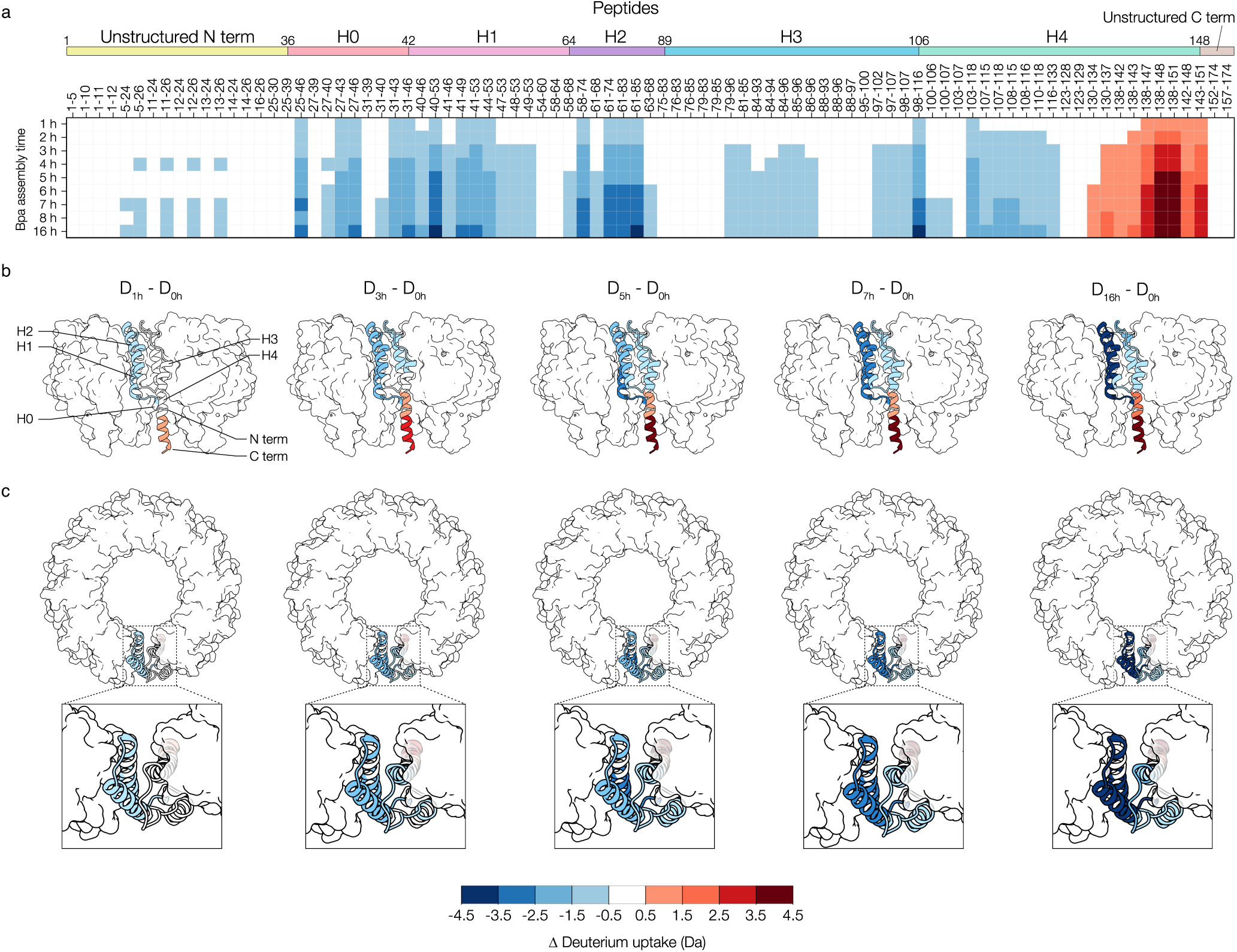
Pulsed HDX-MS reveals the time-dependent evolution of structure during dodecameric Bpa assembly. (**a**) Heatmap showing differential deuterium uptake following a 10-second D2O pulse at various time points during Bpa assembly, relative to the onset of the assembly reaction (*t* = 0 h). Bpa secondary structure elements and their corresponding residue numbers are denoted along the top of the heatmap; Side (**b**) and top views (**c**) of Bpa, shown as surface representations, with one subunit displayed as a cartoon (PDB: 5LFJ)^13^. Differential deuterium uptake is mapped onto the cartoon subunit. Structural figure panels were generated using ChimeraX 1.90^50^.

To further dissect the kinetics of Bpa temperature-induced assembly, we selected peptides exhibiting clear bimodal isotopic distributions. The observed bimodality is indicative of a minimum of two slowly interconverting conformations which can be quantified to extract interconversion rates. We quantified the relative populations of protected (closed – red traces) and exposed (open – green traces) conformations at each incubation time point by fitting a pair of Gaussian functions to the obtained profiles (Fig. 3a). We modelled the time-dependent changes in the fractional population of each conformation using a first-order exponential model for all peptides displaying bimodality (Fig. 3b). This analysis yielded spatially-resolved estimates of the interconversion rate constant (*k*HDX) (Fig. 3c). Across all peptides analyzed, *k*HDX values averaged 0.23 ± 0.06 h^-1^, in excellent agreement with the oligomerization rate determined using SEC (Fig. 1f, Fig. 3c, grey bar). Together, these data define the structural and kinetic features that govern Bpa self-assembly and demonstrate that formation of the dodecamer requires dissociation of cold-stabilized lower-order intermediates that adopt a distinct packing interface.

**Figure 3.**
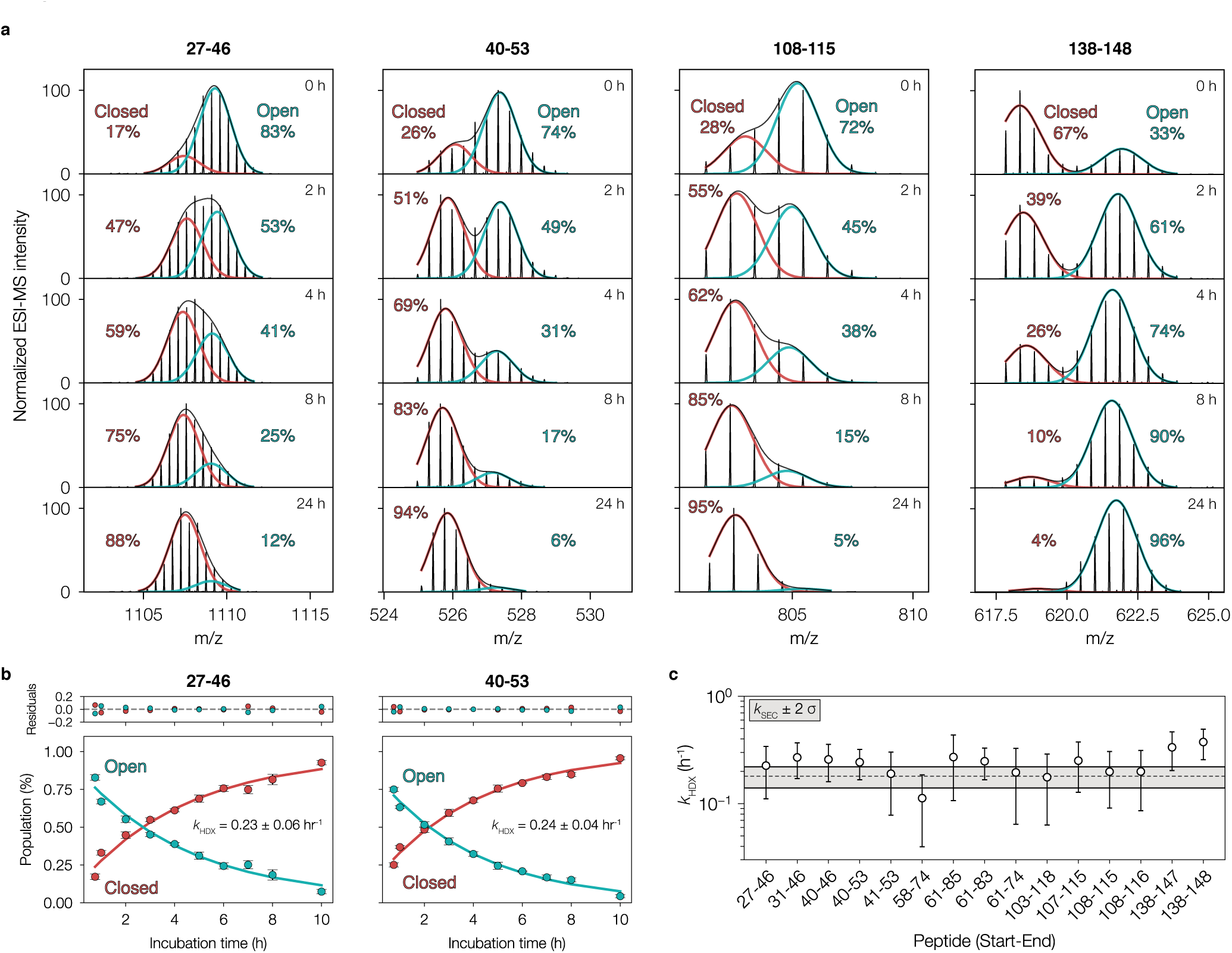
Bpa oligomerization interfaces exhibit asymmetric isotopic distributions during assembly, as revealed by pulsed HDX-MS. (**a**) A pair of Gaussian functions were fit to intensity profiles from peptides displaying asymmetric isotopic distributions and subsequently integrated. Closed (low HDX) and open (high HDX) conformations and their fractional populations are denoted in the raw mass spectra. The displayed peptides are representative of others across the sequence but were selected to maximize sequence coverage; (**b**) Assembly time-dependent changes in the fractional populations of the open and closed conformations for two representative peptides were analyzed using a single-exponential decay model to extract the association rate constant (k). Error bars represent the standard deviation across technical replicates (n = 3); and (**c**) Association rate constants *k* extracted from all peptides exhibiting EX1 kinetics. Error bars represent the standard deviation across three technical replicates. The hatched horizontal grey line denotes the association rate constant determined from SEC measurements, with the shaded region indicating ±2σ. Although the data were not globally fit, the similar *k* values across peptides suggest they report on a common underlying process, namely Bpa assembly.

### Characterization of WT Bpa assembly using methyl-TROSY NMR

In order to examine how temperature affects the structure and oligomerization of Bpa, we turned to NMR experiments, exploiting a methyl-transverse relaxation optimized spectroscopy (TROSY)^34^ effect that enhances spectral sensitivity and resolution for large proteins^34–36^. Methyl-TROSY is particularly effective when paired with selective isotopic labelling of methyl groups (^13^CH3) from Ile, Leu, Val, and Met residues in an otherwise uniformly deuterated ([U-^2^H]) molecule to produce [U-^2^H; Ileδ1-^13^CH3; Leu,Val- ^13^CH3/^12^CD3; Met-^13^CH3]-labeled protein, where only one isopropyl methyl group of Leu and Val is ^13^CH3 labeled in a non-stereospecific manner; hereafter referred to as ILVM-labelling^37^. We recorded ^1^H-^13^C heteronuclear multiple-quantum coherence (HMQC) spectra that benefit from the methyl-TROSY effect^34^ to monitor the temperature-dependent structural changes of Bpa. We obtained high-quality ^1^H-^13^C HMQC spectra at 40 °C and 18.8 T for ILVM-labeled WT Bpa samples that were preincubated at either 4 or 40 °C (Fig. 4a, multi-contour peaks). Prolonged incubation at 40 °C led to the emergence of a distinct set of peaks compared to those obtained from preincubating Bpa at 4 °C, resulting from structural changes to Bpa (Fig. 4a) and corresponding to separate conformers at low and high temperatures. Moreover, the time dependence of the interconversion was slow on the NMR chemical shift timescale as separate peaks were observed for both species at intermediate temperatures.

**Figure 4.**
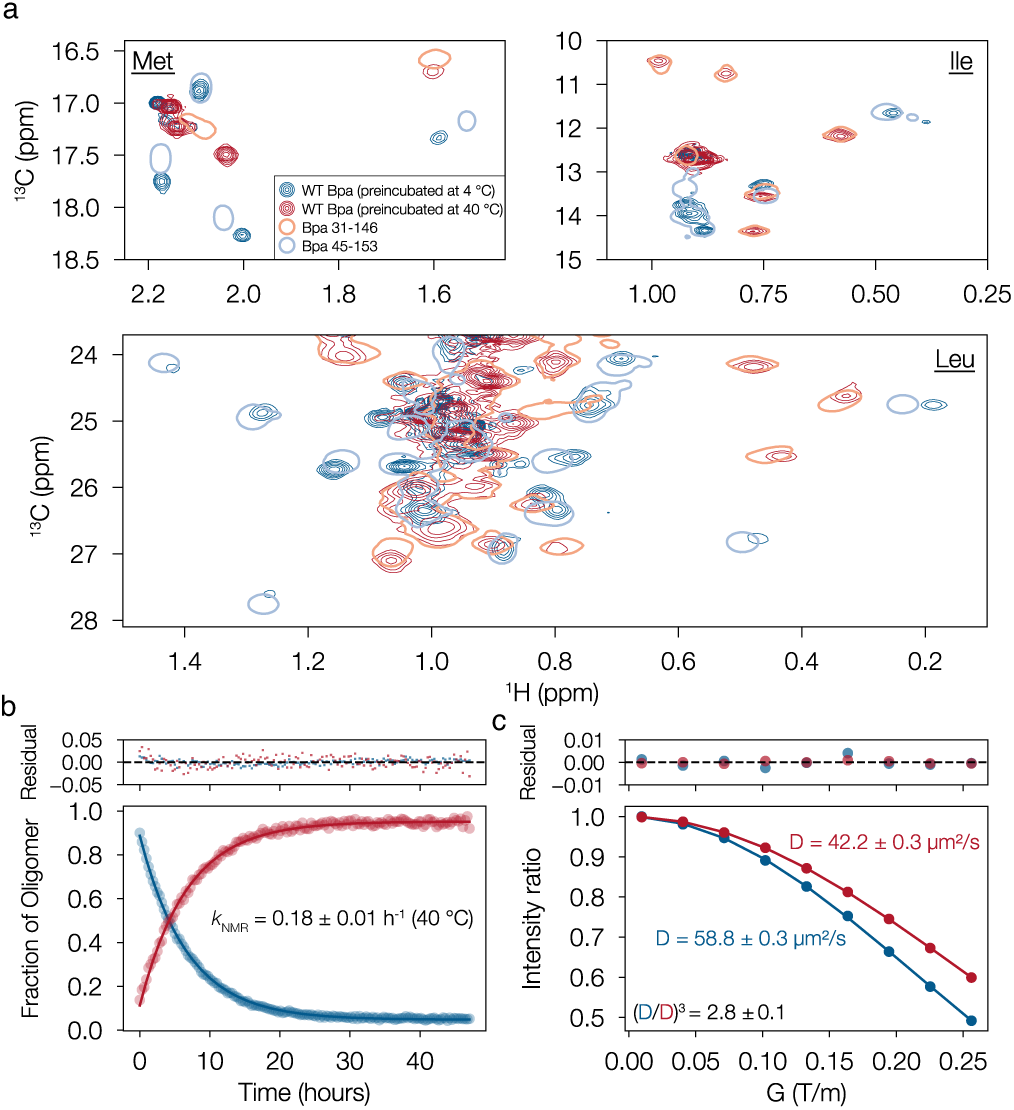
N- and C- terminal truncations stabilize specific oligomeric forms of Bpa. (**a**) ^1^H-^13^C HMQC spectra for ILVM-labeled Bpa WT preincubated at 4 °C (multi-contour blue) or 40 °C (multi-contour red), Bpa 31-146 (single-contour red), and Bpa 45-153 (single-contour blue). All spectra were recorded at 40 °C and 18.8 T; (**b**) Changes in peak intensity for each oligomeric state (tetramer - blue, dodecamer - red) during a 48-hour acquisition. Data were fit assuming a first-order exponential model. Residuals are displayed in the plot above; and (**c**) Intensity ratios of selected peaks (see Supplemental Methods) as a function of gradient strength (G) recorded in a pulsed-field gradient diffusion experiment^51^ (circles). Intensities for a given value of G are normalized to their values obtained with the lowest value of G used in the experiments and fit (solid lines) to extract diffusion coefficients (D). Diffusion coefficients of Bpa WT for each oligomeric state (tetramer - blue, dodecamer - red) are indicated.

To monitor the kinetics and the nature of the oligomeric states involved in this interconversion, we recorded ^1^H-^13^C HMQC spectra over a 48-hour time course, starting from a sample pre-incubated at 4 °C for one week (*t* = 0; Fig. S4). Spectra were acquired every 19 minutes so that the evolution of peak intensities could be quantified rigorously over the period of conversion. For well-resolved peaks, we tracked the time course of NMR signals unique to each of the two oligomeric species and fit the data using a first-order exponential model, yielding an interconversion rate *k*NMR of 0.18 ± 0.01 h^-1^ at 40 °C (Fig. 4b) that agrees well with interconversion rates of 0.18 ± 0.02 h^-1^ and 0.23 ± 0.06 h^-1^ both measured at 37 °C using the SEC (Fig. 1f) and HDX-MS (Fig. 3c) approaches, respectively. To better understand the nature of the oligomeric states involved in this interconversion, we recorded ^1^H-^13^C HMQC-based pulsed-field gradient diffusion experiments focusing on the methionine region over a 48-hour time course during which Bpa was incubated at 40 °C starting from 4 °C at *t* = 0, to infer the size of each species. The resulting diffusion coefficients were 42.2 ± 0.3 µm^2^ s^-1^ for the larger oligomeric species of Bpa present at the end of the incubation period and 58.8 ± 0.3 µm^2^ s^-1^ for the smaller species observed only at the beginning of the time course (Fig. 4c). Diffusion coefficients scale inversely with the cube root of molecular weight^38–40^. The cube of the ratio between the diffusion coefficients for each oligomeric state of Bpa yielded a value of 2.8 ± 0.1, consistent with the presence of dodecameric and tetrameric Bpa species, as supported by native MS and native CD-MS (Fig. 1).

To simplify further NMR studies, we designed constructs where the temperature-dependent oligomerization behavior of Bpa was eliminated. AlphaFold3 (Fig. S5) and our HDX-MS data (Fig. S6a) indicate that the N and C termini of WT Bpa are disordered. Further, X-ray crystallography studies of Bpa demonstrated that N-terminal truncations significantly impact Bpa oligomerization^12^. For example, a Bpa construct spanning residues P44 to Q153, Bpa 44-153, which lacks the N-terminal helix H0, crystallizes as a tetramer composed of a dimer-of-dimers in which the subunits adopt packing interactions that are fundamentally distinct from those observed in the dodecameric assembly (Fig. S3). In the dodecameric structures of Bpa^12,13,20^, H0 packs between helices H2, H3, and H4, an arrangement that contrasts with the inter-subunit H4-H4 contacts observed in the tetrameric assembly^12^. Therefore, deletion of H0 results in an alternative inter-subunit H4-H4 interaction that favours the tetrameric architecture. Guided by these structures and our HDX-MS data which showed that the C-terminal region of Bpa is integral for the formation of the tetrameric species (Fig. S3), but not for the dodecamer, we introduced N- and C-terminal truncations to stabilize specific oligomeric states of Bpa. The methyl-TROSY NMR spectrum of Bpa 31-146 closely resembled that of Bpa WT incubated at 40 °C (i.e. dodecameric Bpa), while the spectrum of Bpa 45-153 was similar to that of Bpa WT incubated at 4 °C (i.e. tetrameric Bpa) (Fig. 4a). Thus, exclusion of H0 and most of the C terminal residues involved in tetramer formation (i.e., the Bpa 31-146 construct) shifts the oligomeric equilibrium toward the dodecamer, independent of temperature. Therefore, we selected the Bpa 31-146 construct, which forms a stable dodecameric structure (Fig. S6b), for all subsequent NMR-based experiments.

### The DNA binding domain of human hTRF1 is a non-native substrate of Bpa-20S CP

Although HspR is the only well-characterized native substrate of Bpa, its poor solubility and lack of stability in isolation^2,11^ have hindered detailed structural investigations. Our attempts to study the Bpa-HspR interaction using NMR spectroscopy were unsuccessful due to HspR’s limited stability and the prolonged NMR acquisition times required. Coexpression of Bpa and HspR mitigates some of these issues but precludes titration-based assays and selective isotopic labelling of each protein for NMR-based measurements. To overcome these constraints, we sought to identify a non-native substrate that would be amenable for probing Bpa-substrate interactions. We focused on the DNA-binding domain of <u>h</u>uman telomeric <u>r</u>epeat-binding factor 1, a 53-residue fragment (residues 378-430 of the full-length protein, referred to hereafter as hTRF1, with residue numbering starting at 1 for this construct) that adopts a well-defined three-helix bundle fold with an unfolded fractional population of ∼1% at 35 °C (PDB: 1BA5)^41,42^. hTRF1 is a well-established model system for NMR-based studies of client-chaperone interactions^42–47^. Notably, hTRF1 is also a known client of the Hsp70 chaperone DnaK, which likewise engages HspR^16,42–45^.

As in previous work^48^, we independently expressed and purified the α- and β-subunits of *Mtb* 20S CP, which were subsequently reconstituted *in vitro*. We also heterologously expressed and purified WT *Mtb* Bpa and hTRF1 and demonstrated using a protease assay that hTRF1 is efficiently degraded in the presence of both WT Bpa and WT 20S CP (Fig. S7). In contrast, a Bpa variant with a Y173A substitution within the C-terminal GQYL motif together with WT *Mtb* 20S CP failed to support hTRF1 degradation (Fig. S8). Similarly, hTRF1 was not degraded when WT Bpa was paired with a catalytically inactive 20S CP variant (βT1A) (Fig. S9). These results establish that hTRF1 degradation is dependent on both Bpa and a functional 20S CP.

### NMR mapping and quantification of the Bpa-hTRF1 interaction

We fully assigned the Met (3 residues; Fig. 5a), Ile (6 residues Fig. 5b), and Leu/Val (17 Leu and 5 Val residues Fig. 5c) regions of a 2D ^1^H-^13^C HMQC spectrum of ILVM-labeled Bpa 31-146, hereafter referred to as Bpa, recorded at 40 °C and 18.8 T. Stereospecific assignments for Leu and Val methyl groups were not obtained. The spectrum of ILVM-labelled Bpa is of outstanding quality, characterized by numerous well-resolved and isolated peaks. We sought to map the Bpa residues responsible for binding hTRF1. Upon the addition of [U-^2^H] hTRF1 to ILVM-labelled Bpa, new peaks appeared for residues near the N terminus of helix H1 and the C terminus of H4 (Fig. 5a-c, light and dark purple contours). Both of these regions are positioned near the base of the Bpa ring^12,13,20^ and precede its unstructured C terminus (e.g., V41, V47, M48, L97, L129, I133, L137, M142) (Fig. 5d). Residues showing chemical shift perturbations (CSPs) clustered into two main regions that are consistent with sites of interaction: (1) the tightly-packed interface between adjacent Bpa subunits (I50, L93, L97, L100, L129), and (2) the more solvent-exposed lower ring (V41, V47, M48, I133, L137, M142). The solvent-exposed residues form hydrophobic bands within the interior of the ring structure and are accessible for possible substrate engagement (Fig. 5d, Fig. S10). The inner ring displays a negatively-charged patch located near the middle of H4 (e.g., E127) and a positively-charged patch at the C terminus of H4 (e.g., R145) (Fig. 5d, Fig. S10). These areas are proximal to V47, M48, L129, and M142, that showed CSPs, suggesting that electrostatic interactions also contribute to hTRF1 recognition^2^. Notably, the hydrophobic residues at the C-terminal portion of helix H4 that exhibit CSPs and are accessible for substrate binding in the dodecameric Bpa are either fully or partially buried at the inter-subunit interface in the tetrameric construct (Fig. S11), suggesting that their accessibility is regulated by the Bpa oligomerization state.

**Figure 5.**
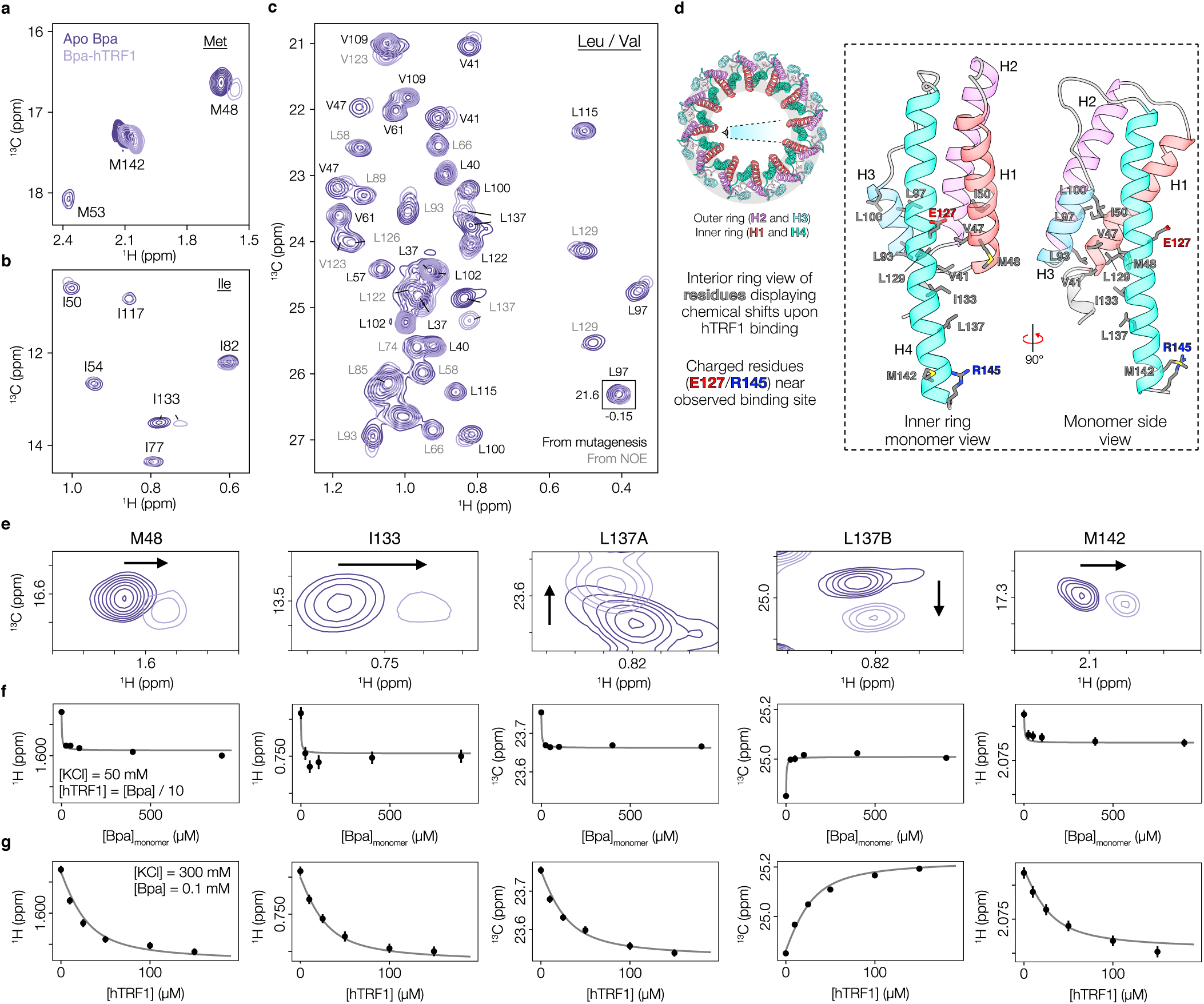
hTRF1 interacts with the C-terminal residues in the lower inner ring of dodecameric Bpa. Assigned (**a**) Met, (**b**) Ile, and (**c**) Leu/Val regions of a two-dimensional 1H-^13^C HMQC spectrum for ILVM-labeled Bpa recorded at 40 °C and 18.8 T. Peaks depicted with dark purple contours are derived from apo Bpa and those with light purple contours from a complex of Bpa and [U-^2^H] hTRF1 in molar ratios of 100 µM (subunit concentration of Bpa): 150 µM (hTRF1) in the presence of 300 mM KCl; (**d**) Top view of the Bpa complex highlighting the inner and outer ring architecture. The inset shows a single Bpa subunit in cartoon representation highlighting residues, shown as grey sticks, that display CSPs upon hTRF1 binding (PDB: 5LFJ)^13^. The sole charged residues proximal to these residues are shown in red and blue; (**e**) Isolated regions of the ^1^H-^13^C HMQC spectrum for residues showing major chemical shift changes upon hTRF1 addition; (**f**) Titration of [U-^2^H] hTRF1 with ILVM-labelled Bpa at 50 mM KCl, monitoring 1H or ^13^C chemical shift changes; and (**g**) Titration of ILVM-labelled Bpa with [U-^2^H] hTRF1 at 300 mM KCl, monitoring ^1^H or ^13^C chemical shift changes.

To quantify the strength of the Bpa-hTRF1 interaction we performed NMR titrations using [U-^2^H] hTRF1 and ILVM-labelled Bpa at a pair of salt concentrations (50 mM or 300 mM KCl) to evaluate the effects of ionic strength on the association (Fig. 5e-g). For experiments performed at 50 mM KCl, we found that the addition of equimolar amounts of hTRF1 to Bpa (subunit concentration) led to the formation of visible protein aggregates and, concomitantly, an increase in solution turbidity. Therefore, we reduced the hTRF1 concentration by 10-fold and worked at a [Bpa]:[hTRF1] ratio of 10:1, starting at concentrations of 900 μM (subunit concentration) and 90 μM for Bpa and hTRF1, respectively, and then serially diluted the mixture and recorded ^1^H-^13^C HMQC spectra, tracking changes in the ^1^H or ^13^C chemical shifts (Fig. 5e). In contrast, at 300 mM KCl there was no evidence of protein aggregation or solution turbidity, and a standard titration was performed where the concentration of Bpa was kept constant at 100 μM (subunit concentration) with [hTRF1] varied (Fig. 5f). For residues exhibiting clear peak displacements as a function of added hTRF1 (Fig. 5e-g), the data were fit to a model assuming exchange kinetics (hTRF1 on/off) that are fast on the NMR chemical shift timescale to extract binding affinities and stoichiometries. Notably, fits of <u>C</u>arr-<u>P</u>urcell-<u>M</u>eiboom-<u>G</u>ill (CPMG) relaxation dispersion profiles recorded with a salt concentration of 50 mM and with several different concentration ratios of Bpa and hTRF1 to a two-site exchange model (interconversion between Bpa free and bound with hTRF1) indicate exchange rates greater than 1500 s^-1^ (Fig. S12). The rates obtained are significantly larger than differences in ^13^C chemical shifts between free and bound Bpa (Fig. 5e), validating the assumption of fast exchange. We further assumed that hTRF1 binds to Bpa only when unfolded, as the ^1^H-^13^C HMQC spectrum of ILVM-labelled hTRF1 in the bound form clearly indicates an unfolded ligand (Fig. S13). The equilibrium constants for the hTRF1 folding/unfolding reaction at 50 mM and 300 mM KCl, that are also required for our analyses, were established by recording <u>c</u>hemical <u>e</u>xchange <u>s</u>aturation transfer (CEST) experiments^49^ (Fig. S14), which yielded unfolded hTRF1 fractions of ∼0.14 and ∼0.07, respectively.

The titration data were fit to a model in which dodecameric Bpa successively binds *n* copies of hTRF1, independent of salt concentration, with a distinct global microscopic dissociation constant at each [KCl] value (see Supplementary Methods). Using unfolded hTRF1 fractions indicated above, we estimated dissociation constants *K*d for the Bpa-hTRF1 complex of < 0.2 µM at 50 mM KCl (Fig. S15a) and 1.5 ± 0.2 µM at 300 mM KCl (Fig. S15b). A plot of reduced-*χ*^2^ versus *n* exhibited a well-defined global minimum with a binding stoichiometry of three hTRF1 molecules per Bpa dodecamer (Fig. S15c). Notably, when it is assumed that hTRF1 can bind equally well to Bpa as an unfolded or folded substrate an identical *n* value is obtained, with approximately ten-fold higher *K*d values at each salt concentration. Together, these experiments identify the hTRF1 binding interface on Bpa, quantify the model-dependent interaction strength as a function of ionic strength, and establish the stoichiometry of the hTRF1-Bpa complex.

### Disordered substrate recognition is mediated by hydrophobic patches on hTRF1

The identification of the Bpa-binding interface on hTRF1 is complicated by the fact that hTRF1 unfolding and binding are coupled, as the free form of the ligand (i.e. hTRF1) is predominantly folded (see above; Fig. S12), while the bound form is not. To this end, we generated a panel of [U-^15^N] hTRF1 deletion variants that lack the ability to adopt a structured fold. We recorded ^1^H-^15^N HSQC spectra of the variants in the absence and presence of 10-fold monomer equivalents of Bpa 31-146 at 300 mM KCI and 40 °C (Fig. 6a). We obtained high-quality datasets characterized by well-resolved resonances focused on a narrow ^1^H chemical shift window that is expected for unfolded proteins. We obtained complete backbone assignments for the hTRF1 fragments under these conditions (Fig. 6a). Upon titration with Bpa, we measured a marked reduction in signal intensities for a subset of hTRF1 amide resonances (Fig. 6a, b). The attenuated cross-peaks map to two distinct sequence segments: residues ^6^WLW^8^ and ^30^ILL^32^, corresponding to a pair of hydrophobic segments (Fig. 6b) previously identified as functionally relevant in hTRF1-DnaK (client-chaperone) interactions^42^. The selective disappearance of peaks results from extensive line broadening caused by two factors: an increase in the effective rotational correlation time of hTRF1 upon binding to the 155 kDa Bpa 31-146 particle, and enhanced transverse relaxation due to the proximity of hTRF1 amide probes to protons from Bpa. In this case, Bpa was fully protonated, which promotes more efficient relaxation than would be observed with a perdeuterated binding partner. Regions of hTRF1 that are no longer visible in HSQC spectra thus correspond to interaction sites with the Bpa client.

**Figure 6.**
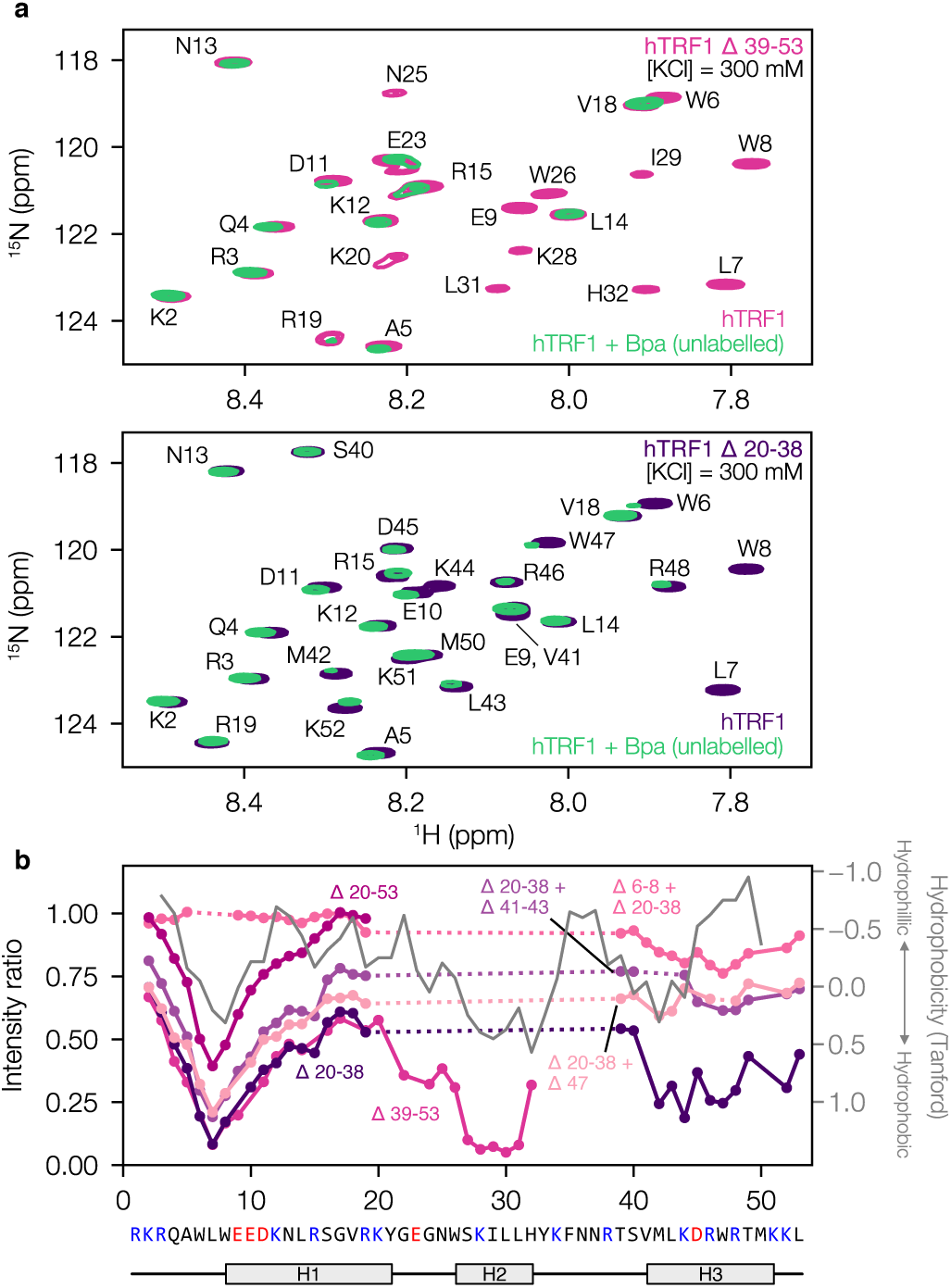
Hydrophobic patches on hTRF1 mediate Bpa binding. (a) Superposition of 1H-^15^N HSQC spectra of [U-^15^N]-labeled hTRF1 in the absence and presence of 10-fold monomer equivalents of unlabelled Bpa 31-146 measured in the presence of 300 mM KCl, 40 °C. Top and bottom spectra were recorded on samples of hTRF1 Δ39-53 and hTRF1 Δ20-38, respectively. (b) Intensity ratios for each hTRF1 peak in the absence and presence of unlabelled Bpa 31-146. The intensity ratios for each hTRF1 construct are colour-coded as indicated. The grey trace indicates the Tanford hydrophobic index^52^ for the hTRF1 sequence.

The hTRF1 deletion variants can also be used to assess whether the affected residues function as independent or cooperative binding elements. Removal of residues 30ILL^32^ (e.g., Δ20-38, Δ20-53, Δ20-38, Δ41-43) failed to eliminate binding near ^6^WLW^8^, suggesting that the two sites act independently. In contrast, deletion of ^6^WLW^8^ abrogated all line broadening effects in the N-terminal region, suggesting its essential role in regulating N-terminal substrate contacts with Bpa. Sequence analysis reveals that both 6WLW^8^ and ^30^ILL^32^ are the most hydrophobic segments of the construct (Fig. 6b – grey trace) and are flanked by polar and charged residues, a motif frequently observed in Bpa substrates and proposed to facilitate substrate pre-orientation via electrostatic steering, followed by hydrophobic engagement at the proteasomal binding interface^2^. Notably, structurally diverse proteins (e.g., FynSH3, drkNSH3, Im7, α-synuclein) that span a broad range of isoelectric points (Fig. S16) did not interact with Bpa as measured by NMR (Fig. S17), underscoring the sequence-encoded specificity of the Bpa-hTRF1 interaction. These findings establish Bpa as a selective reader of short hydrophobic motifs embedded within disordered regions, hallmarks of proteasomal substrates.

## Discussion

Prior cellular, biochemical, and structural studies have established Bpa’s importance within the *Mtb* proteasome system^2,6,7,11–15,20^, yet the mechanisms governing the assembly and substrate recognition by its ATP-independent regulatory particle, Bpa, have remained unresolved. Here, we combine native MS, HDX-MS, and methyl-TROSY NMR spectroscopy, approaches not previously applied to Bpa, with a tractable model substrate to delineate the principles of Bpa oligomerization and substrate engagement.

We show that Bpa exists in a temperature-sensitive equilibrium between dimers, tetramers and dodecamers, and only the fully assembled dodecamer is competent for substrate recognition and delivery to the 20S core particle. HDX-MS, native MS, and CD-MS reveal a cold-stabilized tetrameric species whose protection pattern, particularly at helix H4, is consistent with the inter-subunit packing observed in crystallographic studies of a truncated Bpa variant identified as a cold-stabilized tetrameric state that matches the assembly previously captured in crystallographic studies of a truncated Bpa variant^12^. These H4-H4 interactions must be disrupted to permit formation of the functional dodecamer. We further establish hTRF1 as a structurally tractable, non-native substrate of Bpa and use it to elucidate interactions that are key for substrate engagement. Our experiments exploiting methyl-TROSY NMR establish that Bpa recognizes two solvent-exposed hydrophobic surfaces on hTRF1, ^6^WLW^8^ and ^30^ILL^32^, which are flanked by polar residues. These motifs interact with two distinct regions on Bpa: a hydrophobic lower-ring surface and a charged interior pocket, revealing a bipartite binding mechanism (Fig. 5d). Quantitative NMR titrations show that Bpa binds up to three hTRF1 molecules per dodecamer with micromolar affinity, modulated by salt concentration, further supporting the relevance of electrostatics.

Collectively, our *in vitro* findings support a model in which temperature acts as a regulatory switch for Bpa activity. It is to be established if this model is applicable *in vivo*. Although the host core temperatures remain relatively constant at 37 °C, *M. tuberculosis* experiences a dramatic shift in proteostatic stress when transitioning from environmental persistence to the intracellular environment of macrophages. *Mtb* can survive for hours in aerosols and for days to weeks on surfaces where metabolic activity is minimal and proteostatic demands are low. Elevated temperatures, such as those encountered during phagocytosis by the host macrophages, may drive dodecamer formation, thereby enabling recruitment and degradation of substrates like HspR, a key transcriptional repressor of heat shock genes (*dnaK, gprE, dnaJ, hspR*)^2^. We propose that Bpa may function as part of a broader stress-sensing mechanism that leads to proteasomal activation and transcriptional reprogramming in response to changing conditions. However, this model remains to be validated in a cellular context, and future experiments using Bpa variants locked in defined oligomeric states will be essential to assess the physiological relevance of temperature-dependent assembly. Small molecules that trap WT Bpa in its tetrameric state and prevent functional dodecamer formation may offer a novel strategy for disrupting the *Mtb* proteasome system.

By establishing the structural basis of Bpa oligomerization and revealing how it engages substrates via discrete hydrophobic motifs, our work lays the foundation for understanding proteasomal selectivity in *M. tuberculosis*. The tools and constructs developed here provide a blueprint for dissecting regulated proteolysis in bacterial systems. More broadly, this thermosensitive assembly mechanism may represent a general strategy by which bacteria couple protease activation to environmental change.

## Acknowledgements

B.T.V.D. acknowledges support from a Graduate Tuition Scholarship from the University of Guelph. Financial support was provided by Canadian Institutes of Health Research Project Grant PJT451412 to S.V. and a Foundation Grant FND-503573 to L.E.K., and by a Discovery Grant from the Natural sciences and Engineering Council of Canada RGPIN-2021-02843 to S.V and 024-03872 to L.E.K. MS data were recorded at the Mass Spectrometry Facility of the Advanced Analysis Centre, University of Guelph. We thank Dr. Dyanne Brewer (University of Guelph) for assistance with MS measurements. We thank Prof. Ashok Sekhar (Indian Institute of Science Bangalore) for providing backbone resonance assignments of TRF1. We thank Dr. Algirdas Velyvis (University of Guelph) for guidance and helpful discussions.

## Statement of contribution

B.T.V.D., E.R., L.E.K., and S.V. initiated the project; B.T.V.D., E.R., L.E.K., and S.V. designed research; B.T.V.D., E.R., A.H., J.U., L.E.K., and S.V. performed research; B.T.V.D., E.R., A.H., J.U., D.B., K.R., K.G., L.E.K., and S.V. contributed new reagents/analytic tools; B.T.V.D., E.R., A.H., and S.V. analyzed data; B.T.V.D., E.R., L.E.K., and S.V. wrote the paper.

