## Supporting Information for "Assembly and substrate engagement mechanism of the bacterial proteasome activator Bpa"

[1]Department of Cellular and Molecular Biology, University of Guelph, Guelph, Ontario, N1G 2W1, Canada
[2]Department of Molecular Genetics, University of Toronto, Toronto, Ontario, M5S 1A8, Canada
[3]Waters Corporation, Stamford Avenue, Altrincham Road, Wilmslow SK9 4AX, U.K.
[4]Department of Biochemistry, University of Toronto, Toronto, Ontario, M5S 1A8, Canada
[5]Department of Chemistry, University of Toronto, Toronto, Ontario, M5S 1A8, Canada
[6] Program in Molecular Medicine, Hospital for Sick Children, Toronto, Ontario, M5G 1X8, Canada
[7] These authors contributed equally.

This file contains:

Materials and methods
Figure S1: SEC-MALS on apo Bpa at 4 and 37 °C.
Figure S2: Bpa peptide coverage map in HDX-MS experiments.
Figure S3. Crystal structure of tetrameric Bpa (44-153).
Figure S4: NMR-based studies of the kinetics of oligomerization of Bpa
Figure S5: AlphaFold 3 structural prediction and PAE plot for WT Bpa.
Figure S6. Deuterium uptake plots for disordered N- and C-terminal peptides.
Figure S7: WT Bpa, WT 20S CP, hTRF1 discontinuous degradation assay.
Figure S8: Y173A Bpa, WT 20S CP, hTRF1 discontinuous degradation assay.
Figure S9: WT Bpa, T1A 20S CP, hTRF1 discontinuous degradation assay.
Figure S10: WT Bpa surface hydrophobicity and electrostatics.
Figure S11. Bpa residues involved in hTRF1 binding are sequestered in the tetramer.
Figure S12: Multiple quantum CPMG profiles of Bpa.
Figure S13: Methyl-TROSY spectra of hTRF1 in the apo and Bpa-bound states.
Figure S14: CEST-based estimation of the unfolded fraction of hTRF1.
Figure S15: Global fitting of Bpa and hTRF1 $^{13}$C chemical shifts during NMR titrations.
Figure S16. Isoelectric point (pI) values for potential Bpa substrates.
Figure S17: Bpa $_{WT}$ and Bpa $_{31\text{-}146}$ substrate screen.
Table S1: Synapt G2Si instrument parameters for native MS experiments.
Table S2: HDX experimental parameters and data processing results.

### Methods

#### Plasmids and constructs

Codon-optimized genes encoding for *M. tuberculosis* full-length wild-type Bpa (UniProt: *bpa*, P9WKX3), proteasome α-subunit (UniProt: *prcA*, P9WHU1), and proteasome β-subunit excluding the propeptide (UniProt: *prcB*, P9WHT9) were synthesized and inserted into kanamycin-resistant pET24-based vectors (Biobasic, Markham, ON, Canada). All constructs include a N-terminal $His_6$-SUMO affinity tag, excluding the β-subunit which has a C-terminal TEV-SUMO-$His_6$ tag. Point mutations were introduced via the QuikChange method[1]. The DNA binding domain of hTRF1 (residues 378-430 of the full-length protein, Uniprot: P54274) was used as the unnatural substrate of Bpa for NMR experiments. Additionally, truncated hTRF1 constructs were used to probe regions of the substrate implicated in Bpa binding (Δ 6-8; Δ 20-38, Δ 20-38, Δ 20-38; Δ41-43, Δ 20-38; Δ47, Δ39-53). All hTRF1 constructs include a N-terminal $His_6$-SUMO affinity tag.

#### Protein expression and purification

Multiple Bpa constructs (WT, Y173A, 31-146, 45-153), hTRF1 and its truncated variants, proteasome α-subunit, and proteasome β-subunit were expressed and purified. Plasmids were transformed into chemically competent NEB BL21 T7 Express *LysY E. coli* cells. Different growth media were used depending on the type of desired isotope labeling. For the production of natural abundance proteins, cells were grown in lysogeny broth (LB) to an $OD_{600}$ of 0.6-0.8 before induction with 0.2 mM isopropyl β-d-1-thiogalactopyranoside (IPTG). For the production of uniform $^{15}N$-labeled proteins, cells were grown in M9 minimal media, prepared using $H_2O$ milli-Q, with $^{15}NH_4Cl$ as the sole nitrogen source. For the production of perdeuterated proteins, cells were grown in M9 minimal media, prepared using 99% $D_2O$, with $d_7$ glucose as the sole carbon source and, in the case of ILVM labeling, the following precursors were added 1 h before the induction of protein overexpression: 100 mg/L [ε-13C]-methionine (CLM-206-PK; Cambridge Isotope Laboratories), 60 mg/L α-ketobutyric acid (CDLM-7318; Cambridge Isotope Laboratories), and 100 mg/L α-ketoisovaleric acid (CDLM-7317; Cambridge Isotope Laboratories)[2].

All Bpa constructs and 20S CP subunits were expressed overnight at 16 °C while shaking at 180 rpm. Cells were harvested via centrifugation at 5,000 × *g* for 20 minutes for all proteins. Cell pellets were frozen at -20 °C until future purification. All cells were resuspended in 30 mL of lysis buffer (see below) and lysed via sonication (Heat Systems INC Sonicator XL 2020 Ultrasonic Liquid Processor) using a 5-minute program consisting of 10 seconds on and 20 seconds off. Ion metal affinity chromatography (IMAC) buffers for Bpa constructs consisted of 50 mM Tris, 600 mM KCl, supplemented with either 20 (lysis/wash), 50 (strong wash), or 500 (elution) mM imidazole at pH 7.0. The 20S CP IMAC buffers consisted of 50 mM Tris, 300 mM NaCl, supplemented with either 20 (lysis/wash), 50 (strong wash), or 500 (elution) mM imidazole at pH 7.0. Eluted proteins were dialyzed overnight in lysis buffer + 1 mM dithiothreitol (DTT) with the appropriate proteases to cleave the affinity tags. The 20S CP was assembled using the protocol described by Turner *et al.* (2025)[3]. After dialysis and tag cleavage, a second IMAC step was conducted to remove cleaved affinity tags. Size exclusion chromatography (SEC) using either a Superose 6 or Superdex 200 column was performed using an ÄKTA pure™ chromatography system as the final purification step. Purified Bpa constructs were never frozen and stored at 4 °C until further use.

hTRF1 and its truncated variants was transformed, grown, induced, and harvested as described above. hTRF1 cell pellets were resuspended in 30 mL of denaturing lysis buffer and lysed via sonication (Heat Systems INC Sonicator XL 2020 Ultrasonic Liquid Processor) using a 5-minute program consisting of 10 seconds on and 20 seconds off. Initial IMAC was performed under denaturing conditions. Buffers consisted of 50 mM Tris, 6 M Gdn-HCl, supplemented with either 20 (lysis/wash) or 50 (strong wash) mM imidazole at pH 8.0. The elution step was conducted using a buffer containing 50 mM $NaH_2PO_4$, 200 mM KCl, 2 mM EDTA, and 300 mM imidazole at pH 8.0. Endpoints for the strong wash and elution steps were determined via Bradford assay. Eluted proteins were dialyzed overnight in 50 mM $NaH_2PO_4$, 200 mM KCl, 2 mM EDTA, 50 mM imidazole, and 1 mM DTT at pH 8.0 with the appropriate proteases to cleave the affinity tags. After dialysis and tag cleavage, a second IMAC step was conducted to remove the cleaved affinity tags. SEC using a Superdex 75 was performed using an ÄKTA PUR™ chromatography system as the final purification step. Purified hTRF1 was flash frozen and stored at -80 °C until further use.

Protein quality for all stages of IMAC and SEC was assessed using sodium dodecyl sulfate polyacrylamide gel electrophoresis (SDS-PAGE). Final concentrations of all protein stocks were determined spectrophotometrically using a NanoDrop 2000 spectrophotometer. All extinction coefficients for concentration measurements were calculated via ExPASy's ProtParam (https://web.expasy.org/protparam/).

**Discontinuous SDS-PAGE-based protein degradation assays**

Protein degradation assays were performed by incubating Bpa (WT or Y173), 20S CP

(WT or T1A), and hTRF1 at 37 °C over 48 hours. Protein concentration for all reactions consisted of 10 μM Bpa (subunit concentration), 0.14 μM 20S CP, and 10 μM hTRF1. Aliquots were taken at discrete time points and the reaction was quenched by mixing the reaction mixture with Laemlli buffer (4:1), followed by subsequent heat inactivation at ~95 °C for 5 minutes[4]. Visualization of protein degradation was performed using SDS-PAGE. 20 μL of sample was loaded and run into 20% acrylamide gels for all samples to ensure adequate visualization. BlueElf Prestained Protein Marker (5-245 kDa) as a molecular weight standard. Imaging of all gels was conducted on a BioRad ChemiDoc Imaging System (Hercules, CA).

**Size exclusion chromatography – temperature-dependent oligomerization**

Unassembled Bpa (4 °C) at 50 μM (subunit concentration) was incubated at 37 °C and sampled as a function of incubation time to probe oligomeric state. Separate protein stocks were made for each of the time points (0 , 0.5 ,1 , 2 , 3 , 4 , 5 , 6 , 7 , 8 , and 18 h) so as to not remove the same protein stock from the incubator multiple times during the course of the experiment. Protein samples were spun down for 5 minutes at 21,130 × *g* to pellet any aggregated protein, followed immediately by SEC analysis using a Superdex 200 10/300 GL column using an ÄKTA PURE™ chromatography system. The flow rate for all runs was 0.75 mL $min^{-1}$. Each injection consisted of 200 μL of a 50 μM Bpa (subunit concentration) solution. Visualization and fitting of all SEC chromatograms were made using scripts written in house in Python 3.13.5. The P1, P2 and P3 species were fit using three exponentially-modified Gaussians[5]. The integrals of the peaks were then used to derive the fraction population of P1 and P2+P3. The time dependencies of the populations were subsequently fit using a first order exponential model (Equation 1) to extract an association rate, $k$.

$$f(t) = A \cdot e^{-k \cdot t} + B \qquad \text{(Eq. 1)}$$

### SEC-MALS

WT Bpa was subjected to SEC-MALS analysis to determine the molecular weight of all oligomeric species in solution as a function of incubation time at 37 °C using an OMNISEC multi-detector SEC system (Malvern Panalytical, United Kingdom) fitted with OMNISEC RESOLVE and OMNISEC REVEAL modules. 100 µL samples were injected at 2 mg/mL were loaded onto a P3000 Protein SEC column (300×8 mm, Malvern Panalytical) equilibrated in 150 mM NaCl, 50 mM Tris-HCl, pH 7.5 at a flow rate of 1 mL min$^{-1}$. Molecular weight was calculated using light scattering detectors at 90° (right angle light scattering) and 7° (low-angle light scattering). BSA was used as a standard for molecular weight calibration.

### Pulsed hydrogen deuterium mass spectrometry (HDX-MS)

To probe the assembly of Bpa, we performed pulsed HDX-MS after incubating apo WT at 37 °C for defined time intervals. All buffers ($H_2O$ and $D_2O$-based) used for pulsed HDX consisted of 50 mM imidazole, 100 mM KCl, 0.3% $NaN_3$ [w/v] at pH 7.0. The $D_2O$-based exchange buffer consisted of the same components however the pD was adjusted to 6.6 ($pH_{corr}$ 7.0) using the standard electrode method[6]. Unassembled Bpa (32 µM at 4°C) was incubated at 37 °C and periodically sampled before being pulse labelled in $D_2O$-based exchange buffer for 10 seconds. HDX was started via 40-fold dilution of protein into $D_2O$

based exchange buffer (final Bpa monomer concentration of 0.8 μM) resulting in a 97.5% $D_2O$-based buffer. Exchange was quenched via 1:1 (v/v) dilution into a quench solution consisting of 250 mM $NaH_2PO_4$, 3 M Gdn-HCl, and 3 mM n-dodecylphosphocholine at pH 1.52 followed by immediate flash freezing in liquid $N_2$. Final pH after quenching of samples was determined to be 2.5. All quenched aliquots were stored at -80 °C until LC-MS.

Sample handling and reverse phase liquid chromatography was conducted using a Waters ACQUITY™ M-Class UPLC™ with HDX technology. 20 pmol of sample was digested on-line at 15 °C using a nepenthesin-2 immobilized protease column (Affipro, AP-PC777 004, 1 mm × 20 mm). The resulting peptides were trapped for 3 minutes using a BEH™ 778 C18 (1.7 μm, 2.1 mm × 5 mm; Part#: 186003975, Waters) column at a flowrate of 100 μL $min^{-1}$. Peptide separation was achieved on an HSS T3 (1.8 μm, 1.0 × 50 mm; Part#: 780 186003535, Waters) column. Liquid chromatography consisted of an 8-minute linear acetonitrile:$H_2O$ gradient acidified using 0.1% [v/v] formic acid (acetonitrile ramped from 5 – 35%) at 0 °C and at a flowrate of 100 μL $min^{-1}$. The injection port and sample loop were cleaned between each injection to ensure minimal carryover using a solution consisting of 1.5 M Gdn-HCl, 4% (v/v) acetonitrile, 0.8% (v/v) formic acid, and 1.5 mM dodecylphosphocholine.

The LC outflow was then directed to a coupled Waters SYNAPT™ G2Si Q-TOF mass spectrometer equipped with a standard electrospray source operated in positive ion mode with a capillary voltage of +3 kV. The time-of-flight mass analyzer was set to resolution mode with ion mobility enabled and spectra were recorded over the 50 − 2000 m/z range at a scan rate of 0.4 $s^{-1}$. The instrument was externally calibrated using sodium iodide (700008892) over the 50 – 2000 *m/z* range. The instrument was dynamically calibrated during runs by infusing LeuEnk solution (1+, 556.2771 Th) from the LockSpray capillary every 20 seconds at a flow rate of 10 μL $min^{-1}$. The quadrupole was controlled manually and was set to dwell at 300 m/z for all runs.

All HDX-MS experiments were measured in technical triplicate to ensure reproducibility. Peptide mapping was conducted via ProteinLynx Global Server (v3.0.3) using data from undeuterated samples acquired during $MS^E$ experiments performed as detailed previously[7]. Peptide filtering parameters were taken from Sorenson et al. [8]. All spectra for peptides that persisted after filtering were visually inspected to ensure only high-quality spectra were included in the final data set. Undeuterated control samples were prepared and quenched in an identical manner except for being in an $H_2O$-based solutions. All data analysis was conducted using DynamX v3.0 (Waters). All figures were generated using HDgraphiX[9] and ChimeraX 1.10[10].

**HDX-MS isotopic envelope fitting**

All HDX-MS data were analyzed in Python (v3.13.5) using NumPy, pandas, matplotlib, and the lmfit nonlinear least-squares minimization library. Spectra corresponding to individual peptides at different deuterium labeling times were imported using the Waters Data Importer module of UniDec[11] as intensity versus m/z traces. For each peptide, we fit a pair of Gaussian components to the bimodal isotopic distributions, corresponding to protected (closed) and exposed (open) conformational states. The lower m/z distribution for all spectra were denoted as “closed” and the higher m/z distribution was denoted “open”. In the first pass, spectra across all time points were fit simultaneously with minimal constraints. For each peptide, the center, amplitude, and width of each Gaussian function were allowed to vary freely within broad bounds. The purpose of this pass was to determine a consensus set of parameters, in particular the approximate peak positions and widths, that define the isotopic envelope shape for each peptide. This step reduces user bias and ensures that the minimum number of Gaussian functions is used. Parameters from the first pass were then used as starting values for a second round of fits. In this pass, amplitudes were fully optimized for each spectrum, while centers were only allowed to vary within ±1/z of their first-pass values, and width values were allowed to vary within ±20% of the first-pass value. These constraints enforce consistency in the isotopic envelope shape across time points, while still accommodating small shifts in centroid mass and width due to increasing deuterium uptake. Ultimately, each spectrum was modeled using two Gaussian components, one for the closed state and one for the open state, whose amplitudes were independently optimized while centers and widths remained constrained. This strategy improves the robustness of the fitting parameters, prevents overfitting to noise and ensures that measured population changes directly report on protein dynamics in solution. The integral or the closed and open Gaussians were used to calculate fractional populations for each distribution. These populations were then fit using a first-order exponential model (Eq. 1) to extract an association rate *k*.

**Native ESI MS – Temperature dependent oligomerization**

All MS measurements for apo Bpa were performed in positive-ion mode on a Waters SYNAPT G2Si instrument equipped with a nano ESI source. The time-of-flight mass analyzer was set to resolution mode and recorded spectra over the 400-8000 *m/z* range at a scan rate of 1 $s^{-1}$. The quadrupole transmission profile was operated in automatic mode. All other instrument parameters are listed in Table S1. All LC-MS grade reagents were obtained from Fischer Scientific (Hampton, NH). LC-MS grade ammonium acetate (cat. no. 14267) and water (cat. no. w74) were used to prepare native MS buffers. Reagent grade 2,2-difluoroethylamine (DFEA) (cat. no. D4758) was obtained from TCI America (Portland, OR) and added to the ammonium acetate solution to ensure proper buffering within the capillary during native MS experiments[12]. The final pH of the ammonium acetate + DFEA was adjusted at room temperature using LC-MS grade ammonium hydroxide (60046886) and formic acid (A11750). Bpa was buffer exchanged into 100 mM ammonium acetate at pH 7.0 using a 10k molecular weight cutoff Amicon centrifugal concentrator (3 buffer exchange cycles). After buffer exchange, the protein stock (20 µM – Bpa subunit) was centrifuged and filtered using a 0.2 µm syringe filter device. After filtering, Bpa was diluted to a final concentration of 10 µM using 0.4 M ammonium acetate and 1 M DFEA at pH 7.0. The final concentrations of ammonium acetate and DFEA within the sample were 100 mM and 20 mM, respectively. The Bpa protein stock was then incubated at 37 °C overnight and periodically sampled to monitor the oligomeric state of Bpa as a function of assembly time. At discrete time points ranging from 0 – 24 h, 10 µL of Bpa was sampled and immediately loaded into a borosilicate capillary pulled to a fine tip using a P-1000 micropipette puller (Sutter Instruments). Capillary voltage was applied via a platinum wire in contact with the sample inserted through the open end of the capillary. All specific instrument parameters are listed in Table S1. Deconvolution of data collected using the Synapt G2Si instrument was performed using UniDec[11]. All figures were generated using scripts written in house in Python 3.13.5.

### Charge detection mass spectrometry

Apo Bpa was diluted into 10 mM ammonium acetate solution (Invitrogen, AM9070G) to a concentration of 20 µM (Bpa subunit). A 50 µL aliquot of apo Bpa was buffer exchanged three times with 10 mM ammonium acetate solution (400 µL each time) using Amicon ultra centrifugal filters with a 10 kDa molecular weight cut-off (Merck, UFC501008). Buffer-exchanged samples were then incubated overnight at 37 °C. The incubated samples were diluted to a final concentration of 1 µM (Bpa monomer) prior to charge detection mass spectrometry (CDMS) analysis. Mass analysis was performed using a prototype charge detection mass spectrometer with an electrostatic linear ion trap (ELIT), based on the instrument architecture designed by the Jarrold group at Indiana University, which has been described previously[13]. 5 µL of each sample was loaded into separate glass emitters (5 µm internal diameter) and ions were generated by positive-ion-mode static nanoelectrospray ionization (nanoESI) using a home-built source. Ions were trapped in the ELIT for 100 ms and the frequency and amplitude information were converted to m/z and z values, respectively, and ultimately mass (m/z × z). These data were subsequently binned to generate the corresponding spectra (histograms). The spectra were collected until a minimum of approximately 5000 ions were recorded within the mass range of 0-1 MDa. Signal processing and data visualization were performed using software developed in-house. Visualization of CDMS data was performed using in house scripts in Python 3.12.4.

### Nuclear magnetic resonance (NMR) spectroscopy

All NMR experiments were recorded using one of the following spectrometers: Bruker AVANCE III HD 14.1 T, Varian INOVA 14.1 T, or Bruker AVANCE III HD 18.8 T, all equipped with cryogenically-cooled, triple-resonance probes. All datasets were processed with the NMRPipe software package[14]. $^{1}$H-$^{13}$C HMQC[14] and $^{1}$H-$^{15}$N HSQC spectra were recorded using standard pulse schemes[15].

Diffusion experiments for Bpa were recorded using a previously-published pulse sequence[16] that generates $^{15}$N-edited spectra; in the present case where $^{13}$C-edited spectra are recorded, $^{15}$N pulses are replaced by $^{13}$C pulses. Dephasing and rephasing pulsed-field gradients are applied as bipolar pairs along the y-axis to minimize the effect of convection, with each pair of duration 3 ms, and strengths up to 0.26 T/m. The diffusion delay was set to 0.3 s.

The assignment of methyl groups in $Bpa_{31-146}$ was initially carried out by site-directed mutagenesis where leucine or valine residues were singularly replaced by an isoleucine, and isoleucine or methionine residues were singularly replaced by a leucine. This strategy led to the identification of those assignments labeled in black in Figure 5a-c. All other assignments, labeled in grey, were obtained using three types of NOESY experiments: 1) 3D HMQC-NOESY ($^{1}$H[$t_1$] → $^{13}$C[$t_2$] → mixing time → $^{1}$H[$t_3$]) with a mixing time of 250 ms, recording 40 and 72 complex increments, respectively, for the $^{1}$H ($t_{1,max}$ = 17.9 ms) and $^{13}$C ($t_{2,max}$ = 19.9 ms) indirect dimensions, for a total duration of 48 hours; 2) 3D $F_1$-$^{13}$C, $F_2$-$^{13}$C HMQC-NOESY-HMQC ($^{13}$C[$t_1$] → mixing time → $^{13}$C[$t_2$] → $^{1}$H[$t_3$]) with a mixing time of 50 ms, recording 66 complex increments for both indirect dimensions ($t_{1,2max}$ = 18.2 ms), for a total duration of 33 hours; 3) a second 3D $F_1$-$^{13}$C, $F_2$-$^{13}$C HMQC-

NOESY-HMQC with a mixing time of 250 ms, recording 72 complex increments for both indirect dimensions ($t_{1,2max}$ = 19.9 ms), for a total duration of 44 hours.

Methyl $^{13}C$ CEST experiments were recorded for hTRF1 at [KCl] of 50 mM and 300 mM using the DANTE selective saturation scheme[17,18]. A pair of experiments with $^{13}C$ weak $B_1$ fields fields of 10 and 20 Hz were acquired for each sample, while the position of the $B_1$ field was swept in 41 increments over the spectral window of the DANTE scheme (4 ppm).

Methyl CPMG experiments were acquired for ILVM labeled $Bpa_{31-146}$ in the presence of perdeuterated hTRF1 using the TROSY CPMG pulse sequence[19]. The relaxation of the TROSY component of the {$^{13}C$,$^{1}H$} multiple quantum coherence was measured during a fixed CPMG delay of 20 ms with the application of a variable number of $^{13}C$ refocussing pulses (field strength of 17.2 kHz) with $\nu_{CPMG}$ values ranging from 50 Hz to 2 kHz (14 values plus 3 duplicates). The non-TROSY rapidly decaying components of the {$^{13}C$,$^{1}H$} multiple quantum coherence were suppressed by applying a purge element prior to the CPMG period, as previously described[19].

**Titration of the Bpa-hTRF1 interaction by NMR**

As described in the text, two different titration series were recorded, one at 50 mM KCl and the other at 300 mM KCl to evaluate the role of electrostatics in the Bpa-hTRF1 interaction. Experiments at low [KCl] were recorded using a Bpa monomer:hTRF1 ratio of 10:1, with starting [Bpa] and [hTRF1] of 900 µM (subunit concentration) and 90 µM, respectively. $^{1}H$-$^{13}C$ HMQC experiments were recorded for successive dilutions of this sample ([Bpa subunit] = {900 μM, 400 μM, 100 μM, 50 μM, 25 μM} = [hTRF1] × 10), as well as for the reference sample with Bpa alone. A second titration series was recorded for the 300 mM KCl sample using a standard titration where [Bpa] was kept constant at 100 μM (subunit concentration) and [hTRF1] was varied ({0 μM, 10 μM, 25 μM, 50 μM, 100 μM, 150 μM}).

Profiles of chemical shift changes from the two titration series were fit together to a model in which dodecameric Bpa successively binds single copies of unfolded hTRF1 until *n* copies are bound, with a constant global microscopic dissociation constant $K_d$ (i.e., no cooperativity) for each binding event:

$$B_{12}T_{i-1}^{u} + T^{u} \rightleftarrows B_{12}T_{i}^{u} \quad , K_{eq}^{i} = K_D \cdot i/(n-i+1). \qquad \text{(Eq. 2)}$$

In Eq. 2, subscript *i* indicates the number of bound hTRF1 molecules with $1 \leq i < n$, $B_{12}$ and $B_{12}T_i^u$ denote the dodecameric Bpa species and the Bpa dodecamer-hTRF1 complex comprised of *i* copies of the unfolded form of hTRF1, respectively, and $T^u$ denotes the unfolded hTRF1 ligand. The folding/unfolding equilibrium of hTRF1 is given by

$$T^{f} \rightleftarrows T^{u}, \qquad \text{(Eq. 3)}$$

where $T^u/(T^f+T^u) = f^u$ is 0.14 and 0.07 at 50 mM and 300 mM KCl, respectively, as established by CEST measurements of isolated hTRF1 under conditions that are identical to those used for the titrations (Figure S9).

Note that in Eq. (2) the equilibrium constants for each binding event, $K^i_{eq}$, are related to the factor $i/(n-i+1)$ that accounts for the multiplicity of ways in which ligand can be added to $B_{12}T_{i-1}^{u}$ ($n-i+1$) and subtracted from $B_{12}T_{i}^{u}$ ($i$). Profiles at both [KCl] were fit together using Eqs. 2 and 3, with *n* treated as a global parameter, i.e. assumed to be salt-independent, while separate dissociation constant $K_d$ values are fit, one for each [KCl].

In the above analysis it is assumed that Bpa can only bind to unfolded hTRF1 ($T^u$). Alternatively, if Bpa binds to both folded ($T^f$) and unfolded hTRF1 with the same affinity, the same best-fit value of $n$ = 3 is obtained, however the $K_d$ values increase by *ca.* one order of magnitude. Thus, while the $K_d$ values are model dependent, the value of *n* is less sensitive to the mechanism of binding.

### Supplementary figures

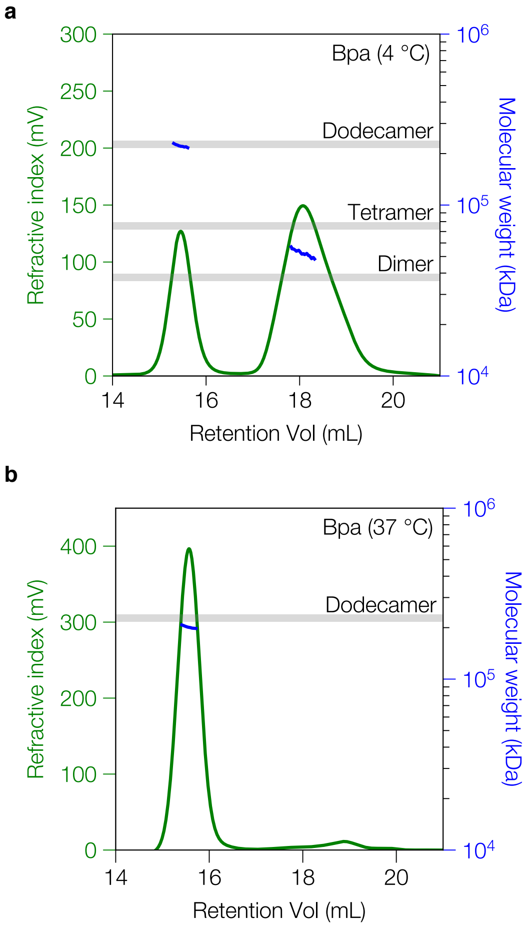


**Figure S1. SEC-MALS analysis of apo Bpa at 4 and 37 °C.** (**a**) Bpa $_{WT}$ incubated at 4 °C. Two distinct peaks are observed, a well-resolved peak corresponding to dodecameric Bpa and a broad peak corresponding to trimeric Bpa. The grey bar indicates the expected molecular weight ± 5%. (**b**) Bpa $_{WT}$ incubated at 37 °C and analyzed at 20 °C. A single predominant peak corresponding to dodecameric Bpa is observed. The left y-axis and green trace represent the refractive index signal, which reflects protein concentration, while the right y-axis and blue trace indicate the estimated molecular weight. Grey bars denote ±5% of the expected molecular weight for each indicated oligomeric species. An OMNISEC multi-detector SEC system (Malvern Panalytical, United Kingdom) fitted with an OMNISEC RESOLVE and OMNISEC REVEAL modules was used.

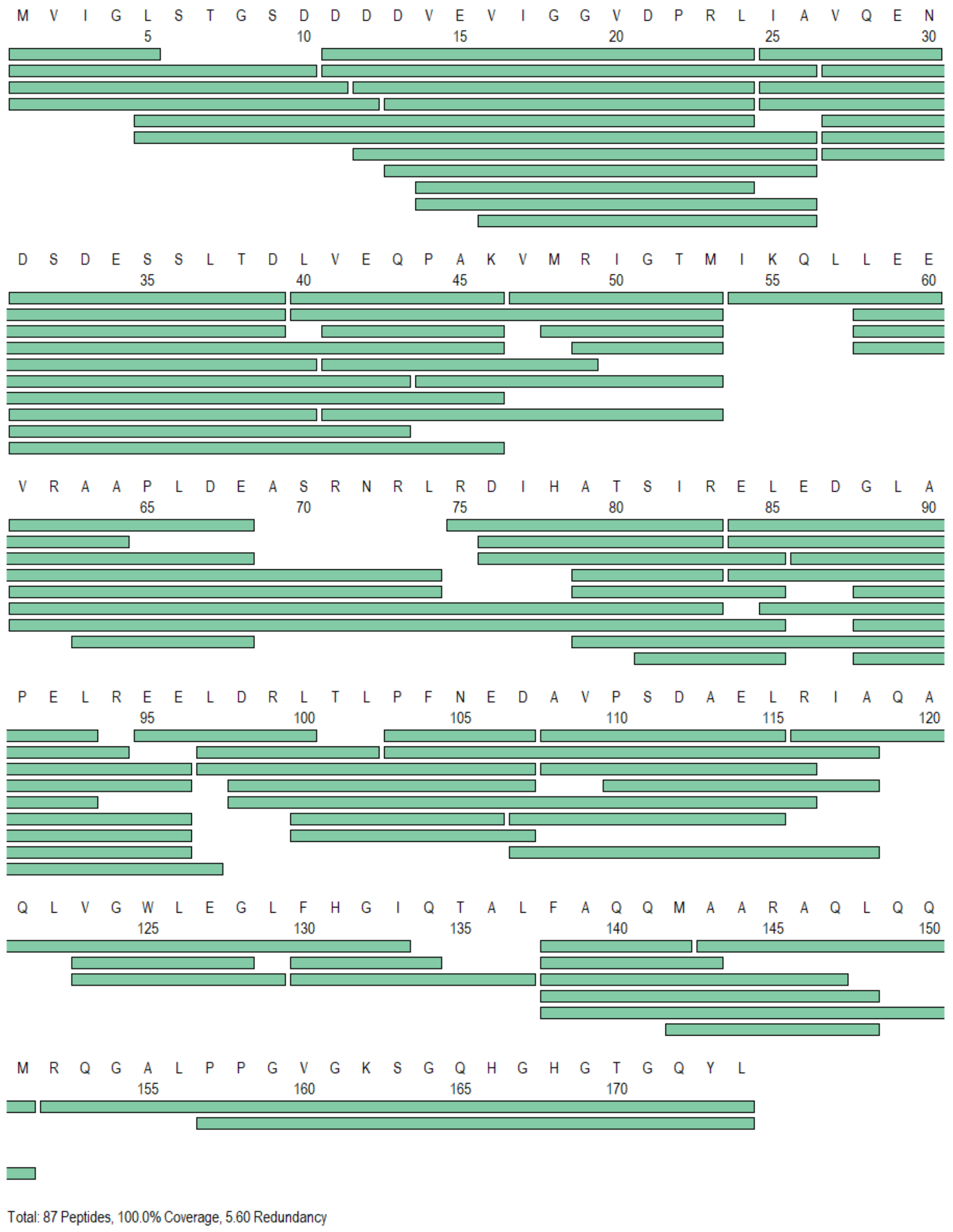


**Figure S2. Bpa peptide coverage map in HDX-MS experiments.** Peptide coverage for Bpa after online digestion using nepenthesin II for pulsed HDX-MS experiments. After filtering using the parameters listed, 87 peptides with quantifiable deuterium uptake remained, corresponding to a sequence coverage of 100% and a peptide redundancy level of 5.60.

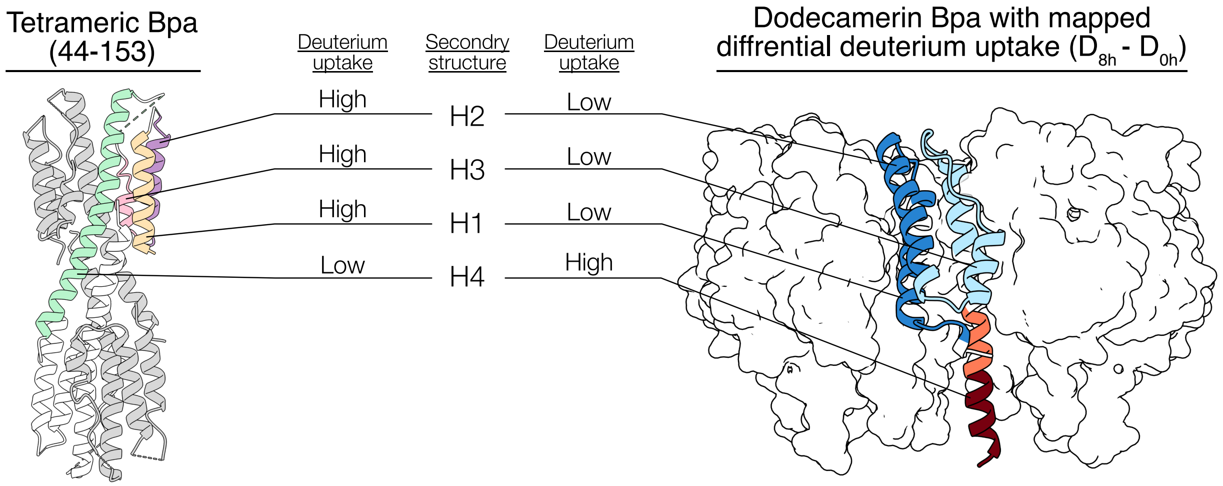


**Figure S3. Structural comparison of tetrameric and dodecameric Bpa assemblies.** Left: Crystal structure of the tetrameric Bpa construct (residues 44–153; PDB: 5IEU)[20], shown as a cartoon with each helix individually colored. Right: Dodecameric Bpa (PDB: 5LFJ)[21], shown as a surface representation, with one subunit displayed as a cartoon and colored according to differential deuterium uptake after 8 hours of assembly (from HDX-MS). Increased deuterium uptake in helix H4 in the dodecamer reflects loss of protection due to conformational rearrangement. In the tetramer, H4 forms a tightly packed inter-subunit interface (green), leading to strong protection. In contrast, helices H1, H2, and H3 are solvent exposed in the tetramer and exchange rapidly, but become buried and protected upon dodecamer formation as they are sequestered between adjacent Bpa subunits.

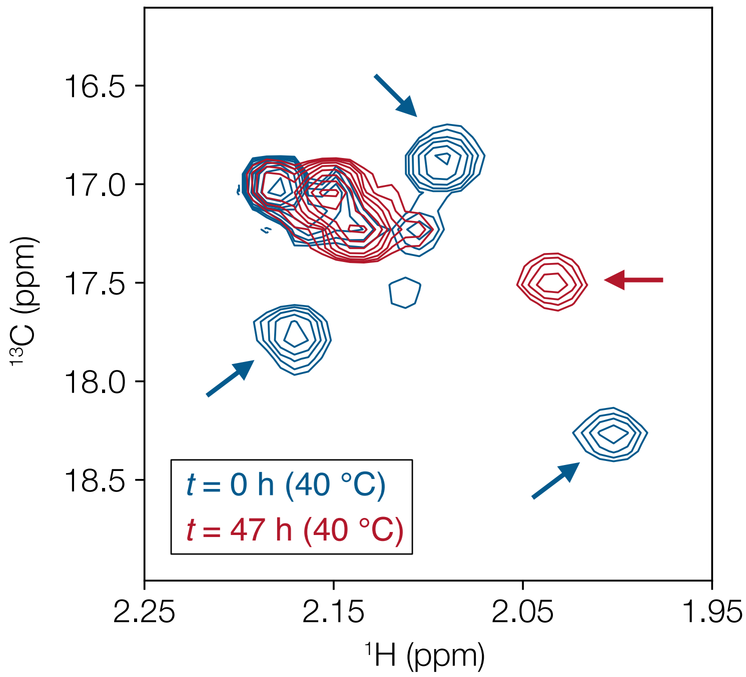


**Figure S4. NMR probes used for the measurement of the kinetics of tetramer-dodecamer interconversion and the establishment of diffusion rates of each species.** We performed a kinetic experiment to follow the disappearance of the low-temperature tetramer and buildup of dodecameric Bpa NMR signals by recording a series of HMQC spectra at 40 °C. Overlay of $^{1}$H-$^{13}$C HMQC spectra of ILVM-labelled Bpa, focusing on the Met region of the data set, at *t* = 0 (immediately after equilibration at 4 °C; blue) and at t = 47 h (red). The arrows indicate which isolated peaks were used in our analyses.

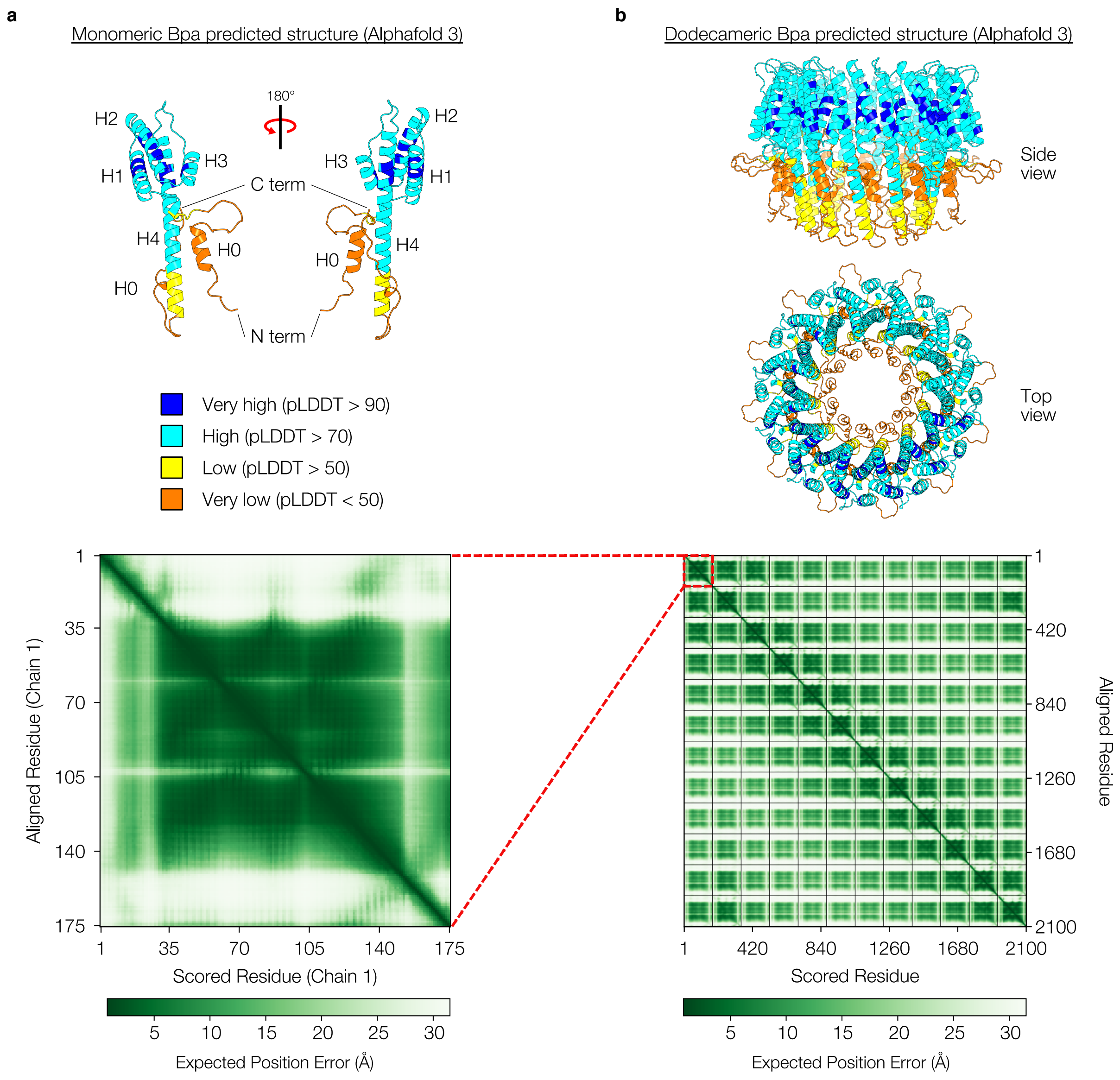


**Figure S5. AlphaFold3 predicts that the N and C termini of Bpa are unstructured.** (a) Cartoon representation of a Bpa monomer predicted by AlphaFold 3 (AF-P9WKX3-F1). The structure is coloured according to the pLDDT score which is indicative of the prediction confidence. All elements of secondary structure and each terminus are labelled; and (b) Predicated aligned error (PAE) plot for the AlphaFold3 predicted structure.

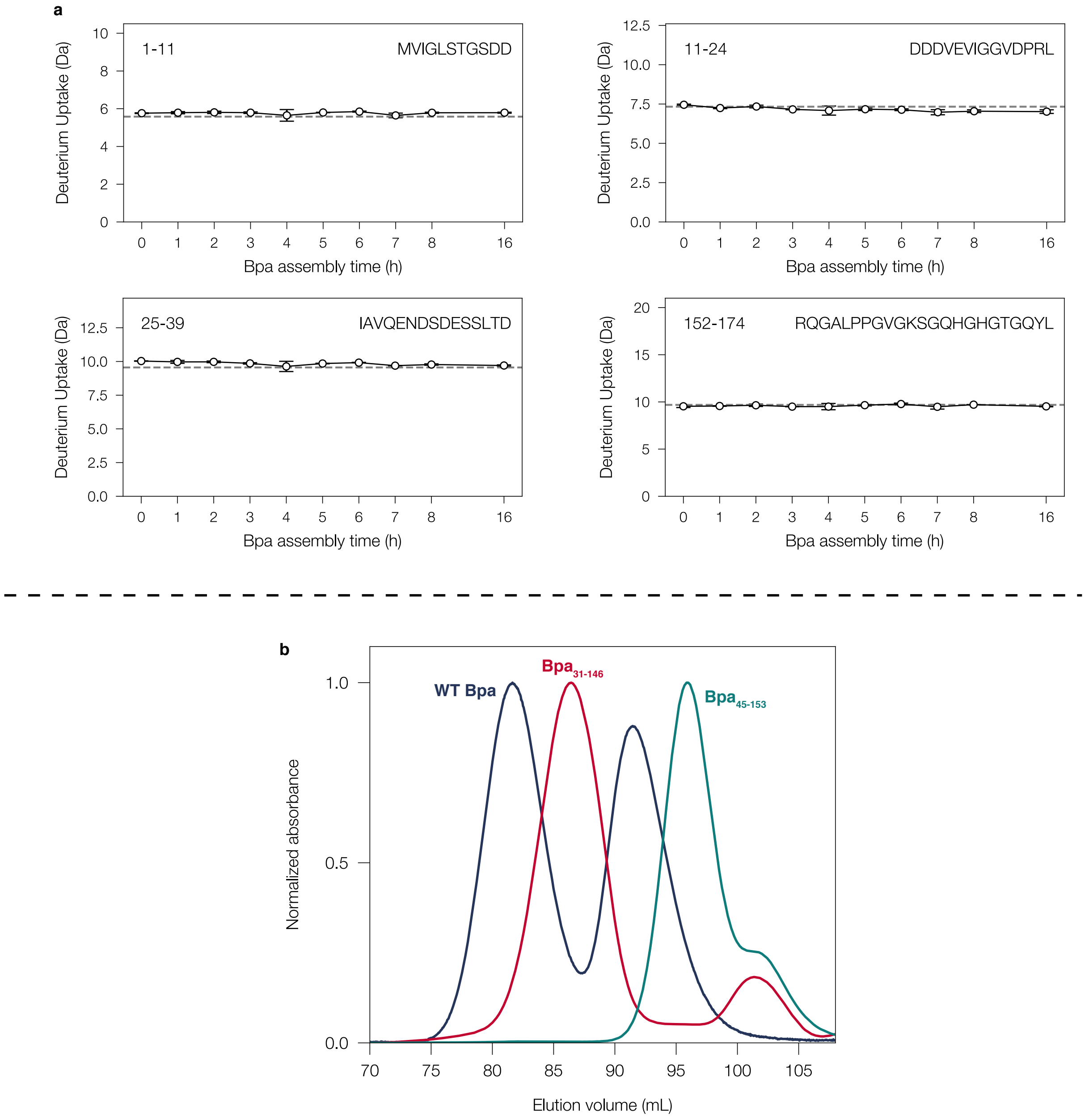


**Figure S6. Deuterium uptake plots for disordered N- and C-terminal peptides.** (a) N- and C-terminal peptides become fully deuterated after a 10-second exposure to $D_2O$ with no observable kinetics as a function of Bpa assembly time indicating unstructured termini. The dashed horizontal line indicates the fully deuterated control. The start and end residue numbers and peptide sequence are indicated in the top left and right corners of each plot, respectively; and (b) SEC profiles of WT Bpa, Bpa $_{31-146}$, and Bpa $_{45-153}$.

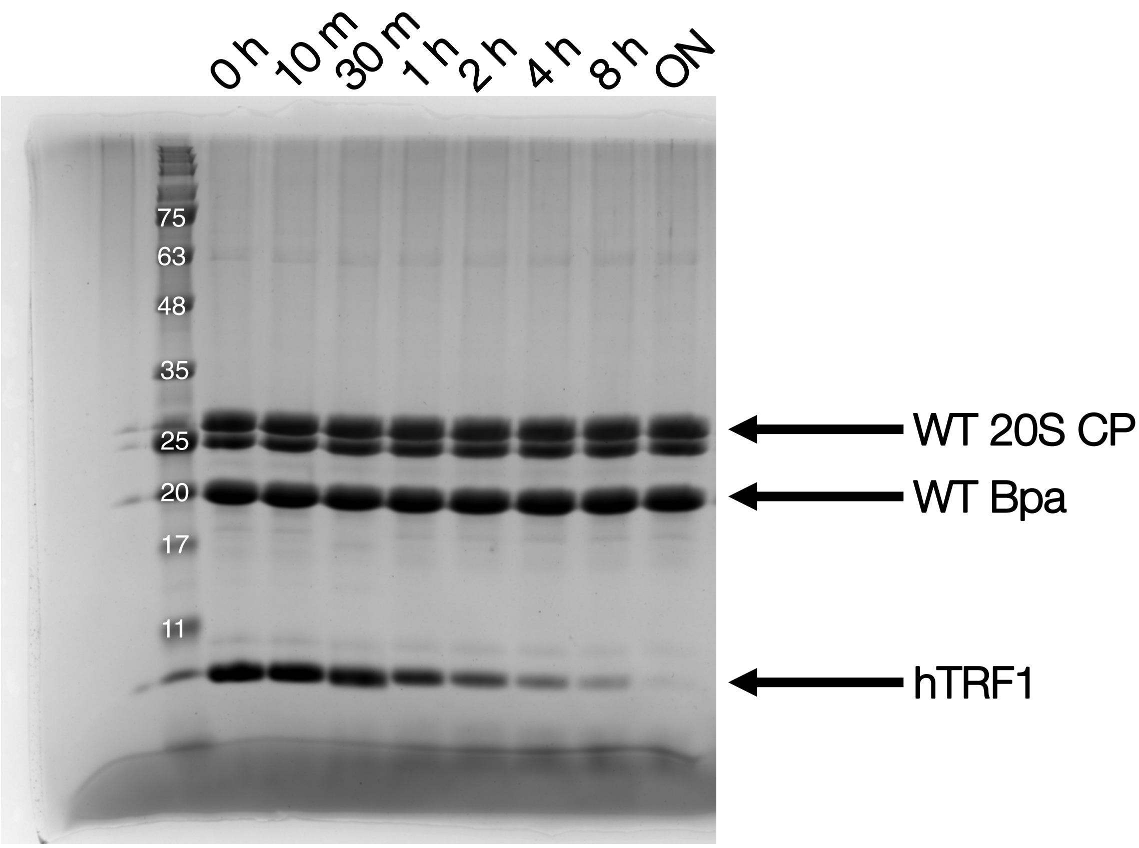


**Figure S7. Bpa-mediated proteasomal degradation of non-native substrate hTRF1.** All components (WT Bpa, WT 20S CP, hTRF1) were combined and incubated at 37 °C for 48 hours. Aliquots were taken at discrete time points and quenched. Reaction protein concentrations were 0.14 μM 20S CP, 0.8 μM Bpa (dodecamer), and 10 μM hTRF1.

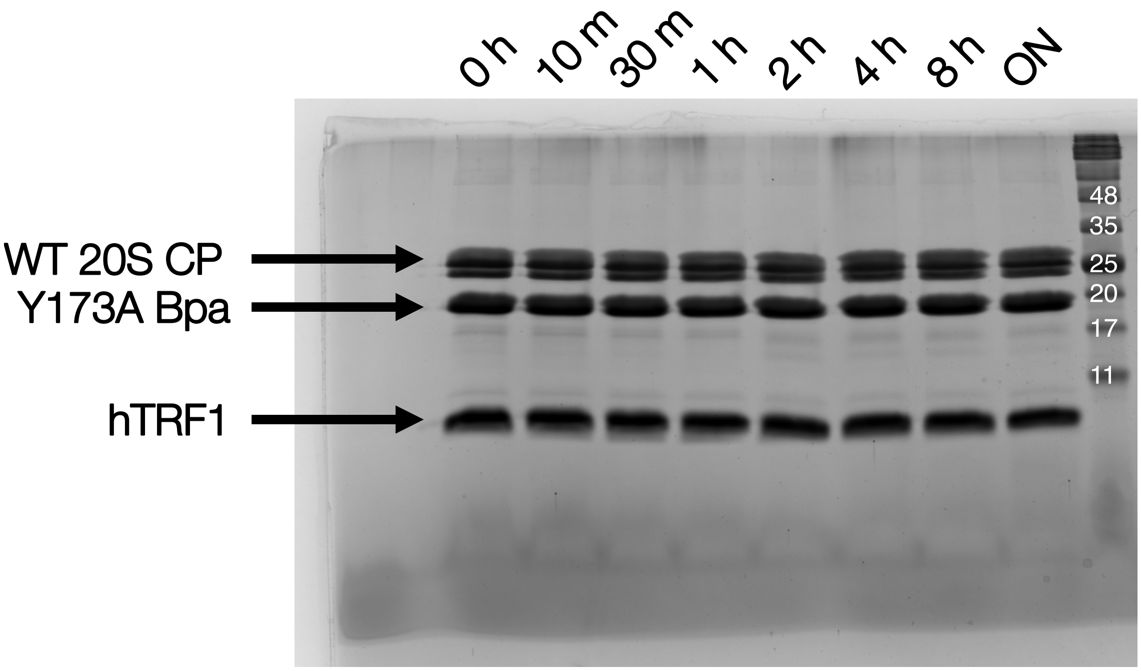


**Figure S8. The Y173A substitution in Bpa abrogates association with 20S CP resulting in inhibited hTRF1 degradation.** Lack of degradation when Y173A Bpa is used implies that GQYL-mediated interaction between WT Bpa-20S (Fig. S5) is required for degradation to proceed. All components (Y173A Bpa, WT 20S CP, hTRF1) were combined and incubated at 37 °C for 48 hours. Aliquots were taken at discrete time points and quenched. Reaction protein concentrations were 0.14 μM 20S CP, 0.8 μM Bpa (dodecamer), and 10 μM hTRF1.

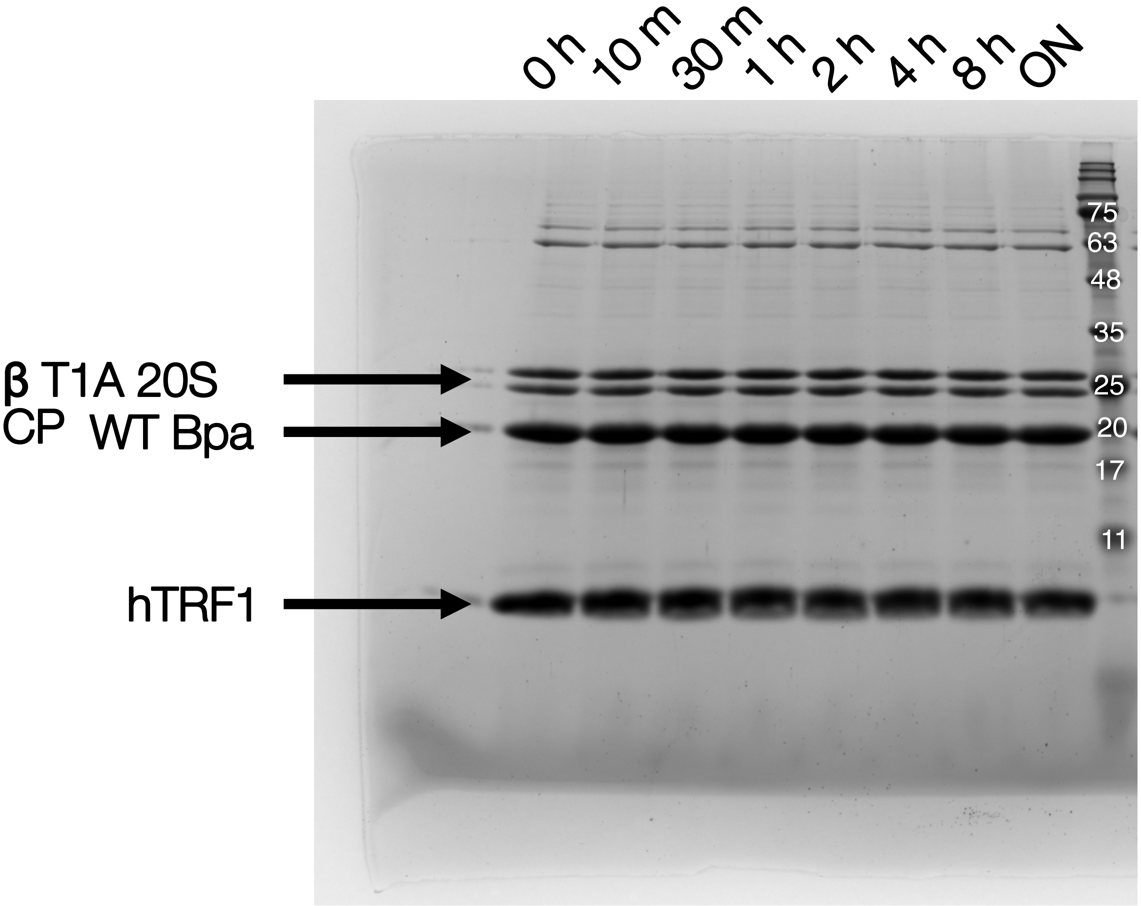


**Figure S9. hTRF1 is not degraded by the catalytically-inactive 20S CP.** All components (WT Bpa, βT1A 20S CP, hTRF1) were combined and incubated at 37 °C for 48 hours. Aliquots were taken at discrete time points (0 h – 48 h) and quenched. Reaction protein concentrations were 0.14 μM 20S CP, 0.8 μM Bpa (dodecamer), and 10 μM hTRF1.

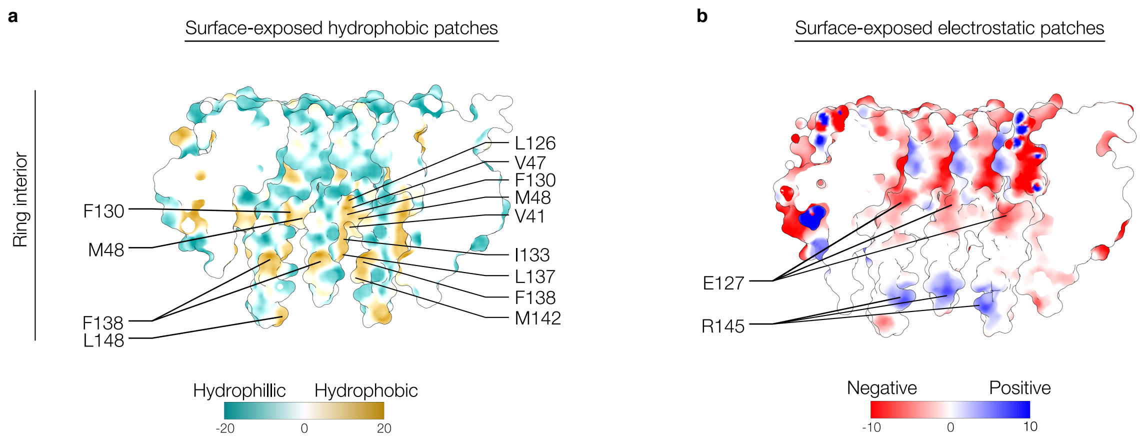


**Figure S10. Bpa $_{WT}$ surface hydrophobicity and electrostatics.** (**a**) Surface-exposed and sequestered hydrophobic patches on the inner-ring of Bpa are shown. Residues contributing to hydrophobicity are labelled and indicated by black lines. (**b**) Surface-exposed and sequestered charged patches for the inner ring of Bpa are shown. Residues contributing to electrostatics are labelled and indicated by black lines. PDB ID: 5LFJ from Bolten *et al.* 2016[21].

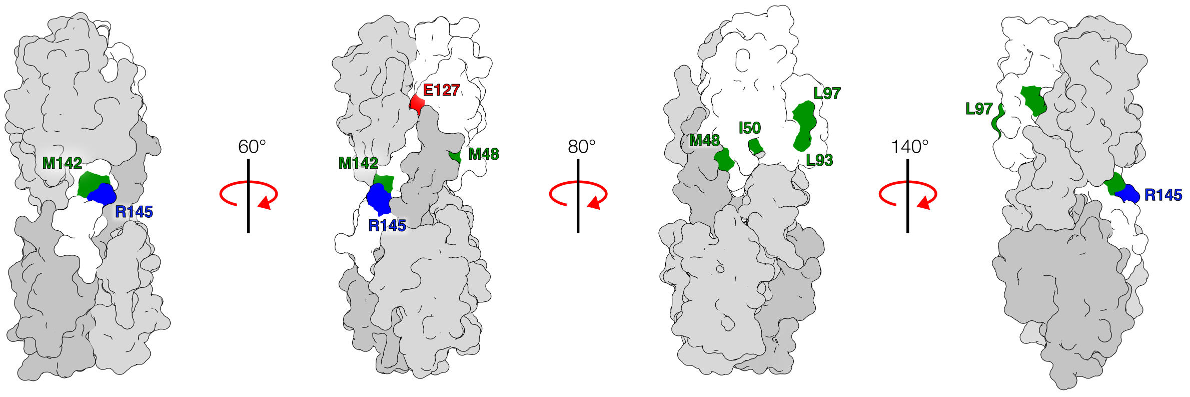


**Figure S11. Bpa residues involved in hTRF1 binding are sequestered in the tetrameric form.** The tetrameric Bpa structure is shown as a surface representation with one monomer highlighted in white (PDB: 5IEU). Green patches correspond to surface-exposed residues that also display chemical shift changes upon addition of hTRF1. Red and blue patches denote electrostatic regions relevant to substrate engagement in the dodecameric form. All colored residues are labeled.

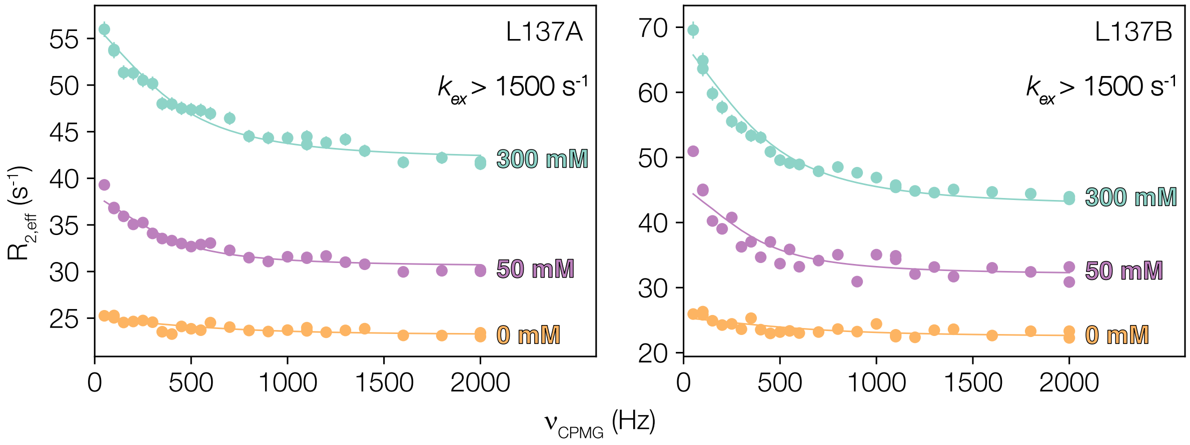


**Figure S12. {$^{13}$C-$^{1}$H} multiple quantum CPMG profiles of Bpa.** CPMG profiles for the two methyl groups of L137 (A, left; B, right) at [Bpa] = 900 μM (subunit concentration) and [hTRF1] = 0 μM (orange), 45 μM (purple), and 90 μM (green). Note that the $^{13}$C nuclei of the methyl groups of L137 are two of the probes used in chemical shift-based fits of titration profiles to establish hTRF1 binding affinities and stoichiometries (Figure 5).

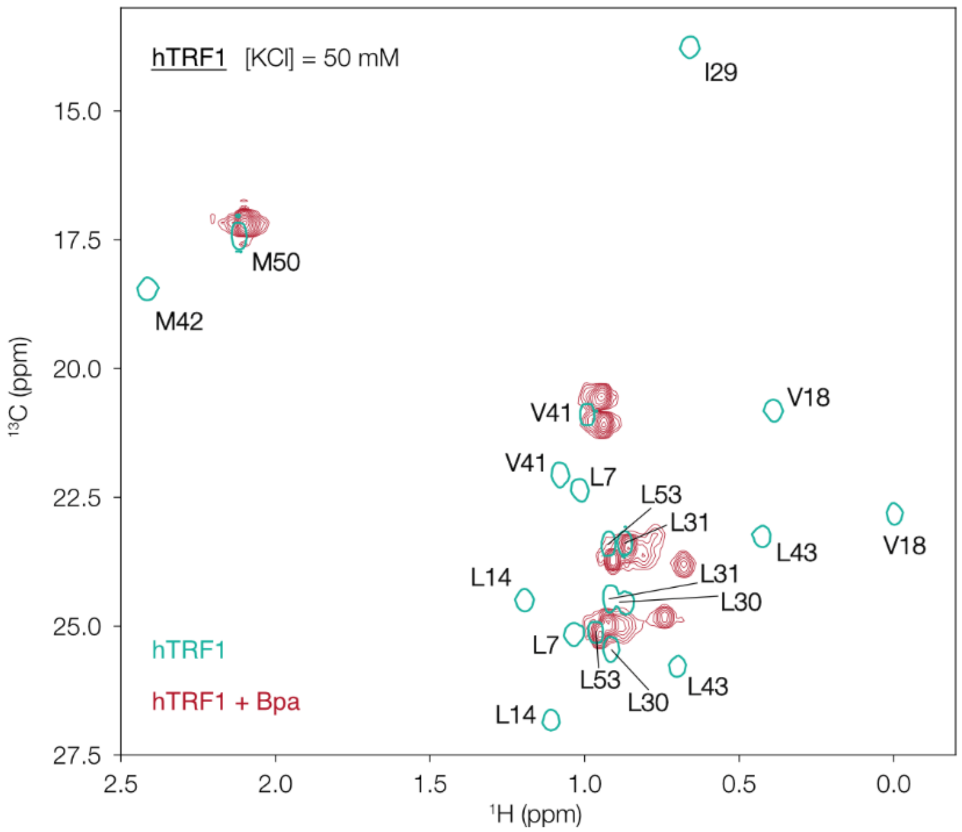


**Figure S13. hTRF1 is unfolded when bound to Bpa.** Superposition of a selected region from $^{1}H$-$^{13}C$ HMQC data sets (40 °C, 600 MHz) recorded on samples of ILVM-labeled hTRF1 (turquoise, single contours) and ILVM-labeled hTRF1 in complex with protonated Bpa (red, multiple contours), 50 mM KCl. Note that the spectra of isolated hTRF1 is well-dispersed, while in the chaperone-bound state hTRF1 is only poorly resolved.

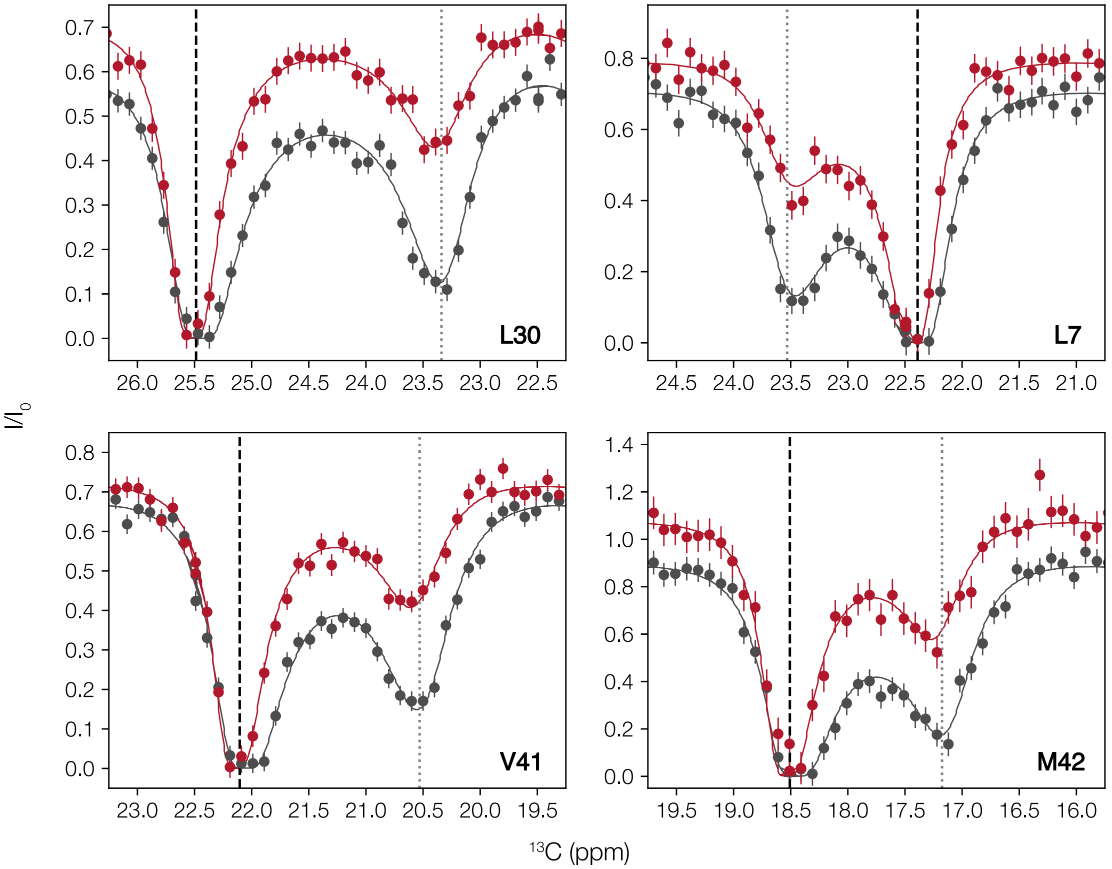


**Figure S14. Estimation of the unfolded fraction of hTRF1 at 50 and 300 mM KCl, 40 °C.** $^{13}$C methyl D-CEST profiles[17,18] for four representative methyl groups (denoted on each spectrum) at [KCl] = 50 mM (grey) and 300 mM (red). The chemical shifts of the folded state are indicated by vertical black dashed lines, while those of the unfolded state are indicated by vertical gray dotted lines. Note that the depth of the dips for the unfolded state at high salt (red) are reduced compared to those at low salt (grey), indicating a stabilization of the folded state at the higher ionic strength. Quantitative analysis of the data using the program Chemex[22] indicates a decrease in the unfolded fraction, $f^U$, from 13.6 ± 0.4% at [KCl] = 50 mM to 7.1 ± 0.3% at [KCl] = 300 mM.

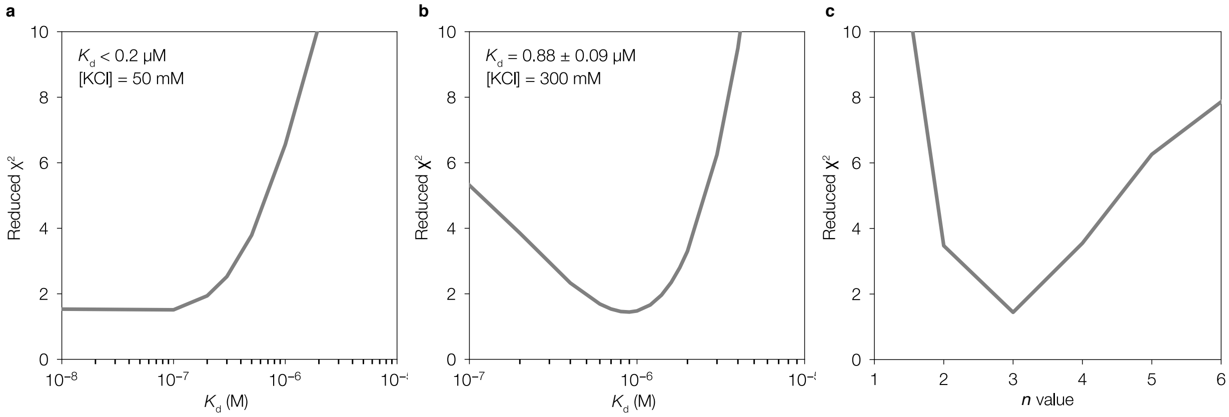


**Figure S15. Analysis of hTRF1 – Bpa binding via fits of titration profiles.** Reduced $\chi^2$ as a function of the microscopic dissociation constant (Eq. 2), $K_d$, for hTRF1 binding at 50 mM (a) and 300 mM (b) KCl. Values of $K_d$ were obtained from joint fits of all the titration data (both salt concentrations), assuming that the number of bound hTRF1 ligands is independent of salt concentration. A binding model where only the unfolded state of hTRF1 associates with Bpa is assumed. (c) Reduced $\chi^2$ as a function of *n*, establishing that three copies of hTRF1 bind to dodecameric Bpa.

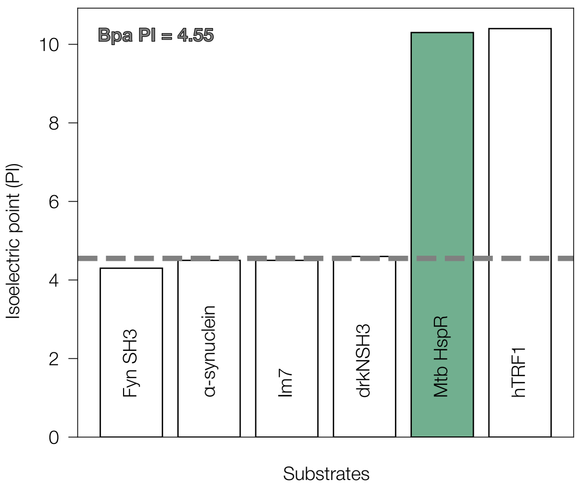


**Figure S16. Isoelectric point (pI) values for different proteins tested as potential Bpa substrates.** Each bar denotes the pI of an experimentally tested or well-established substrate. The green bar denotes HspR, a native substrate of Bpa in *Mtb*. The grey hatched line indicates the pI of Bpa, also shown in the top left corner.

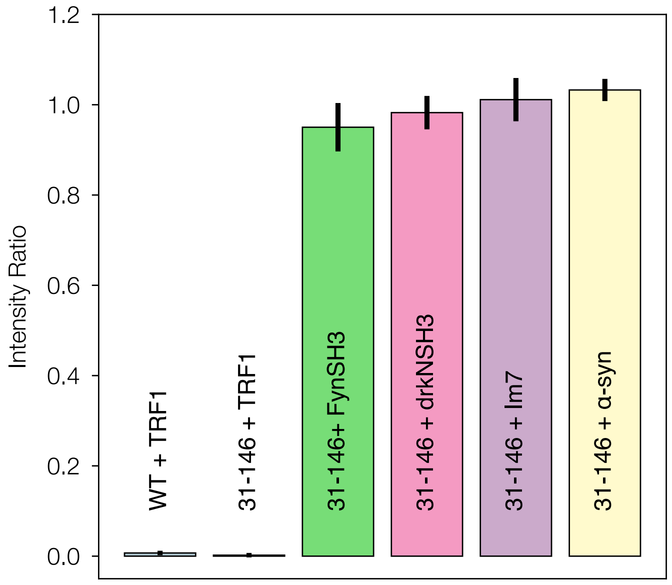


**Figure S17. Bpa $_{WT}$ and Bpa $_{31\text{-}146}$ substrate screen.** NMR peak intensity ratio when Bpa was combined with various substrates. A low peak intensity ratio indicates binding. The Uniprot accession numbers for each substrate are as follows: hTRF1 (P54274, residues 378-430 of the full-length protein), FynSH3 from *Gallus Gallus* (Q05876, residues 85-142, A39V/N53P/V55L mutant[23], drkNSH3 from *Drosophila melanogaster* (Q08012, residues 1-59), Im7 (Q03708), α-syn (P37840).

**Table S1.** Instrument parameters used to record native mass spectra on a Synapt G2Si.

| | |
|---|---|
| **Ion transmission** | |
| ESI capillary voltage | 0.7 -1.0 kV |
| Cone voltage | 80 V |
| Source temperature | 30 °C |
| Desolvation temperature | not applicable |
| Source offset | 80 V |
| **Gas flow** | |
| Cone gas | 50 L/h |
| Desolvation gas | not applicable |
| Nebulizer gas | not applicable |
| Trap gas | 4.0 mL $min^{-1}$ |
| **Collision energy** | |
| Trap collision energy | 0 V |
| Transfer collision energy | 0 V |
| **Trap DC** | |
| Entrance | 1 V |
| Bias | 0 V |
| Trap DC | -2 V |
| Exit | 0 V |
| **IMS DC** | |
| Entrance | -20 V |
| Helium cell DC | 1 V |
| Helium exit | -20 V |
| Bias | 2 V |
| Exit | 20 V |
| **Transfer DC** | |
| Entrance | 5 V |
| Exit | 15 V |
| **Trap and transfer Triwave** | |
| Trap wave velocity | 300 m/s |
| Trap wave height | 0.5 V |
| Transfer wave velocity | 247 m/s |
| Transfer wave height | 0.2 V |
| **IMS Triwave** | |
| IMS wave velocity | 300 m/s |
| IMS wave height | 40 to 20 V over 100% of IMS duty cycle (linear) |
| **Mobility trapping** | |
| Release time | 100 µs |
| Trap height | 15.0 V |
| Extract height | 0 V |
| **Pressures** | |
| Backing | 3.07 mBar |
| Source | $7.61 \times 10^{-3}$ mBar |
| Trap | $1.64 \times 10^{-2}$ mBar |
| Helium | $2.82 \times 10^{-4}$ mBar |
| IMS | $3.63 \times 10^{-4}$ mBar |
| Transfer | $1.46 \times 10^{-2}$ mBar |
| TOF | $1.15\times10^{-6}$ mBar |

**Table S2.** Hydrogen deuterium exchange experimental parameters and resulting statistics.

| | **Apo Bpa** |
|---|---|
| HDX reaction details | Final $D_2O$ concentration [v/v] 97.5%, $pH_{corr}$ 7.0, 37 °C |
| HDX time course (sec) | 1 h, 2 h, 3 h, 4 h, 5 h, 6 h, 7 h, 8 h, 20 h |
| Undeuterated controls | 3 |
| Back-exchange | 33% (value determined by Turner *et al.* 2025)[3] |
| Number of peptides | 147 |
| Sequence coverage | 100% |
| Average peptide length/redundancy | 5.60 |
| Replicates | 3 technical replicates |
| Repeatability | 0.15 Da |
| Significant differences | 0.5 Da |
